# Theoretical analysis of confinement mechanisms for epigenetic modifications of nucleosomes

**DOI:** 10.1101/2022.12.20.521095

**Authors:** Jan Fabio Nickels, Kim Sneppen

## Abstract

Nucleosomes and their modifications often facilitate gene regulation in eukaryotes. Certain genomic regions may obtain alternate epigenetic states through enzymatic reactions forming positive feedback between nucleosome states. How a system of nucleosome states maintains confinement is an open question. Here we explore a family of stochastic dynamic models with combinations of readwrite enzymes. We find that a larger number of intermediate nucleosome states increases both the robustness of linear spreading in models with only local recruitment processes and the degree of bi-stability under conditions with at least one non-local recruitment. Further, supplementing the positive feedback with one negative feedback acting over long distances along the genome enables effective confinement of epigenetic, bistable regions. Our study emphasizes the importance of determining whether each particular read-write enzyme acts only locally or between distant nucleosomes.

## Introduction

Chromatic regions can adopt one of two stable states that they can pass on to the next generation without changes in the underlying DNA sequence (1, 2). Modeling such epigenetic regions revealed essential properties for bistability and heritability like positive feedback of the modifying enzymes, non-local interactions, and (implicit) cooperativity (3). The basic model structure has been successfully modified to answer different questions and to model nucleosome-mediated epigenetics in various organisms, like in, e.g., *S. Pombe* (4–6), *S. Cerevisiae* (7, 8), *Plasmodium falciparum* (9), *Drosophila* (10, 11) and *Arabidopsis thaliana* (12, 13). Often, bistable regions are short and show abrupt boundaries in ChIP-seq profiles of associated histone modifications. Two examples of relatively small bistable regions are the two mating-type regions HML and HMR (around 3 kb) *in budding yeast* (14) and the FLC locus in *Arabidopsis thaliana* (around six kB) (12). Moreover, the simultaneous presence of H3K27me3 marks, associated with silent gene expression, and H3K4me3, associated with active gene expression, characterizes a large subset of important promoters in mouse embryonic stem cells (*mESCs*). Due to the co-occupation of silent and active modifications, these around 1 kb long promoters have been termed ‘bivalent’ (15). However, a recent study suggests that these regions may be bistable, consisting of a heterogeneous population of active or silent cells rather than ‘bivalent’ where both states coexist predominantly on the same locus (10).

Other small, putative bistable regions are heterochromatic islands in fission yeast. These facultative heterochromatic regions have a characteristic size of around 3 kb and are influenced by external factors like iron (16), or caffeine (17). These factors have been shown to regulate the expression of the putative demethylase *Epe1* or *Mst2*.

Although relatively large regions like the mating-type region in S. *pombe* are very well characterized and have been modeled extensively (4–6), mathematical modeling of smaller regions has been scarce. Especially how small regions can remain bistable and heritable despite high direct modifications and noise levels remains elusive.

In addition to more uniformly distributed block-like ChIP-seq profiles, many genomic regions exhibit a bell-shaped pattern with a high central peak and increasingly lower modification levels at the periphery. The coexistence of these distinct profiles suggests different modes of heterochromatic spreading from the nucleation center, even though the same proteins are often found at these different types of loci. Moreover, previous models assume the region of interest to be perfectly isolated from its surroundings. Models, where confinement of histone modifications arises spontaneously due to the underlying rules while accounting for all known properties of the read-write enzymes, have been fragile and assuming variable recruitment strength between different spatial regions (7).

Here we theoretically explore mechanisms that allow small chromosomal regions to be bi-stable and confined. The proposed models are constrained by consistency with previous models and experiments of more extended parts of the genome (3–6). Our analysis pinpoints that more intermediate states increase resistance to noise and enable a more expansive repertoire of functionalities. Furthermore, a combination of positive and one global negative feedback from the heterochromatic state enable confinement of a silent or bistable state even without local barriers. Although the model is inspired by observations of the heterochromatic islands in fission yeast, we believe its underlying mechanisms can be applied to systems in other organisms.

## Results

### Linear-spreading models need multiple recruitment steps for intrinsic confinement in the presence of direct methylation

To examine the necessary conditions/properties of confining histone modifications after being established, we first looked at a well-established linear-spreading model of histone modifications (18, 19) that enables intrinsic confinement. Intrinsic confinement in this context means that the spreading of modifications from a nucleation site is limited to a specific genomic location without the need for explicit boundaries/insulators (e.g., binding sites for transcription factors like CTCF). The authors developed the model based on the experimental observation that H3K9me3 marks propagate gradually and symmetrically from a nucleation center at a synthetically modified *Oct4* locus of pluripotent cells and fibroblasts. Nucleation occurs via HP1*α* that is selectively recruited to the nucleation site upon chemical induction and that associates with H3K9me-specific histone methyltransferases (HMTs) like SETDB1 and SUV39h1/2 (20–23). These HMTs have read-write properties that enable them to bind to H3K9-methylated nucleosomes (the same modification that they catalyze) and to subsequently methylate nearby nucleosomes via allosteric activation of their enzymatic domain (24, 25). Starting from unmodified nucleosomes (H3K9un), this read-write mechanism creates positive feedback, resulting in a symmetric, linear propagation of H3K9me marks from the nucleation site.

The model structure is straightforward, requiring only two parameters, the feedback rate k+ and the demethylation-rate k- (see Methods). For simplicity, the nucleation rate is set to be the same as the feedback (k+). The model is simulated for a total of 50.000 timesteps on a one-dimensional lattice with 257 sites (representing nucleosomes) starting from only H3K9un nucleosomes to ensure that the system settles to a steady state. At each time step, the central nucleosome and each nucleosome adjacent to an already methylated nucleosome are methylated with a probability of k+, and demethylation happens with a probability of k-at every nucleosome in the system. The authors found that the simulations create bell-shaped steady-state profiles of H3K9me confined to a central region up to a ratio of 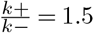 without requiring any explicit barriers/insulators. Confinement is a consequence of weak positive feedback (compared to the demethylation reaction). Thus, the probability of having modified nearest neighbors that can propagate the modification becomes smaller the farther away the modified nucleosomes are from the nucleation site that provides a constant source of newly methylated nucleosomes.

The above model assumes that H3K9-methylation can only occur directly via nucleation sites or by recruitment through neighboring nucleosomes. However, while loss of methylgroups ( H3K9me to H3K9un state transitions) can occur at each nucleosome, direct methyl-group additions (H3K9un to H3K9me) are assumed to be completely absent. We find this assumption overly idealized since the direct collision of histones with HMTs and thus direct methylation at each nucleosome is likely to still occur in addition to feedback reactions, albeit possibly with a much lower probability.

The first question we asked was, what effect a low rate of direct methylation (k+ direct) would have on the confinement of H3K9me marks?

Figure 1 explores the linear-spreading model with the slight modification of adding a direct k+ rate (direct H3K9un to H3K9me transitions). Direct H3K9un to H3K9me transitions here are 50 times less likely than direct H3K9me to H3K9un transitions (fig. 1 B). The model has been simulated for different 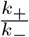 values to obtain different steady-state methylation profiles (Fig. 1 D). Fig. 1 E displays the basal height of these profiles far from the peak as a function of 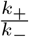. One sees that the intrinsic confinement is entirely lost, even for relatively small values of 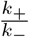 (black line).

**Fig. 1.**
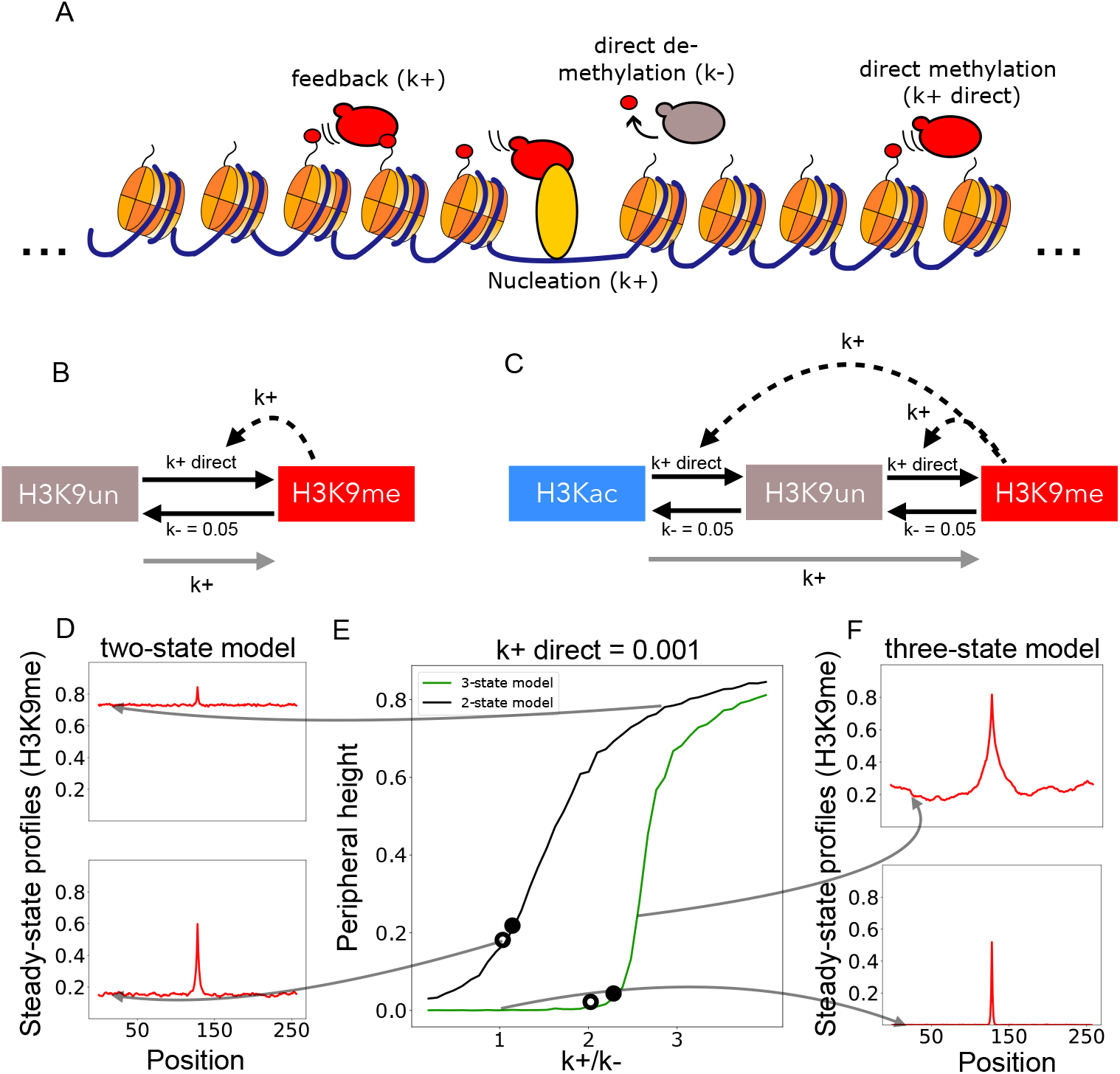
An additional recruitment step from the H3K9me-state increases the effectiveness of confinement in the presence of low direct silencing rates. A) Illustration of all possible interactions of the basic linear spreading model. B) Schematic of a two-state model with local positive feedback from H3K9me-nucleosomes. C) Schematic of a three-state model with local positive feedback from H3K9me-nucleosomes. In both models, direct nucleosome connversions from methylated (H3K9me) to unmodified (H3K9un) are fixed at a rate of *k*— = 0.05 conversion attempts per nucleosome per timestep and direct nucleosome connversions in the opposite direction (H3K9me to H3K9un) are fixed at a rate of 0.001 conversion attempts per nucleosome per timestep. Grey arrows represent nucleation events, where only the central nucleosome is attempted to be converted. For simplicity, feedbacks (curved dashed arrows) and nucleation events are attempted with the same rate k+, whereas the value of k+ is varied in E resulting in a ratio 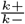. D and F) Example steady-state profiles showing the time-averaged enrichment of H3K9me-modified nucleosomes resulting from simulations corresponding to the 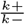 values depicted by the curved arrows. E)Relationship of the height at peripheral position 20 as a function of 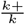 for the two-state and three-state model with fixed *k*+ = 0.001. Black open circles indicate conditions with a peak width of 10 nucleosomes at half-maximal peak height. Black full circles mark conditions with a width of 20 nucleosomes at half peak height. See also Figures S1, S2 and S3.

Subsequently we wondered whether an additional intermediate state subject to feedback from H3K9me would enable intrinsic confinement even with the additional k+ direct reaction. This additional state is justified by the observation that HP1*α* also associates with class II histone deacetylases (HDACs) (24, 26), likely resulting in a similar feedback mechanism.

Including one additional recruitment step in the model (Fig. 1 B) results in perfect spatial confinement for 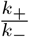 up until a value of ≈ 2.4 with a sharp transition at that value (blue line). We also examined the width of the profiles at half-maximal peak size and found that a broad bounded profile is possible with the three-state model but not with the two-state model (see Figures 1 D and S1).

Additionally, two-state and three-state models with local recruitment and the additional property of long-range nucleation were tested. In these models, similar to (27), nucleosomes are attempted to be converted with a probability of 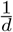 with distance d from the central nucleosome. In this case, additional recruitment steps also help confine methylated regions in the presence of small direct demethylation and acetylation rates (see Fig. S3).

We thus conclude that multi-step recruitment reactions vastly increase the model’s robustness against direct methylation of nucleosomes, effectively enabling the confinement of histone modifications surrounding a nucleation site.

So far, we only investigated linear-spreading models. It is essential to mention that this class of linear-spreading models cannot establish bistable chromosomal regions. Bistable chromatin means that a single locus can be either predominantly marked with histone modifications associated with active gene expression or with modifications associated with a silent gene expression state. Thus, for bistabile systems, intermediate states with a balanced mixture of active and silent nucleosomes need to be unstable. Bistable chromatin has been experimentally demonstrated in different organisms (2, 8, 12). Bistability requires at least one non-local positive feedback (28) and is further strengthened by positive feedback in both directions. Here non-local feedback is mediated by read-write enzymes that are able to bridge between nucleosomes that are distant along the DNA.

A crucial unresolved question is how bistable chromosomal regions are intrinsically and robustly confined by combining local and non-local recruited conversion reactions. Combining several local- (nearest-neighbor only) feedback interactions with a small subset of non-local feedback reactions is further supported by a recent publication about different spreading mechanisms used by yeast ATM and ATR kinases around DNA double-strand breaks (29). We thus now explore possibilities to increase bistability and probe mechanisms that enable intrinsic confinement of histone modifications on small patches of nucleosomes.

### Multiple intermediate states increase robust epigenetics

Models often assume that nucleosomes can only adopt one of two or three states (a silent and an active one) (18, 19, 27). However, histone modifications often come in several degrees. For example, lysine 9 (H3K9me), lysine 36 (H3K36me), lysine 4 (H3K4me), or lysine 27 (H3K27me) of histone three often show multiple methyl groups covalently bound to their amino group. The read-write enzymes catalyzing the addition of these methyl groups can often bind their substrate and modify nearby nucleosomes. PRC2, for example, binds to H3K27me3 histone tails via its RBBP4/7 subunit and catalyzes mono-, di- and trimethylation of H3K27 on nearby nucleosomes (reviewed in (30)). This binding to the nucleosome with the highest degree of methylation and the subsequent sequential addition of methyl groups to nearby nucleosomes creates the possibility for non-processive multistep feedback. Indeed (31) argues for such multi-step recruitment of methylated marks in a model where the opposing reactions are mediated by transcription.

Here, we now explore how multiple recruitment steps influence the behavior of models that also allows for one or several non-local feedback reactions. These explorations extend the previous analysis of (31).

Figure 2 shows the simulated probability distribution of the nucleosome states in a three-state and a 5-state model implemented on a *L* = 10 system. From now on, we generically call the different nucleosome states A*, A, U, S, and S*, representing different modifications- and modification degrees depending on the system in question. E.g., One can think of the S and S* states as representing the PRC2-catalyzed H3K27me and H3K27me2/3 modifications and of the A* and A states as H3K4me and H3K4me2/3 modifications catalyzed by Trithorax (10). Occasionally, we call the group of A* and A nucleosomes active due to their association with active gene expression states without making any claims about a causal connection with transcription itself. Conversely, we denote the combined group of S and S* states as silent nucleosomes due to their correspondence with silent gene expression states. This logic of several modification degrees can be extended to a 7-state model (see Fig. S4)

**Fig. 2.**
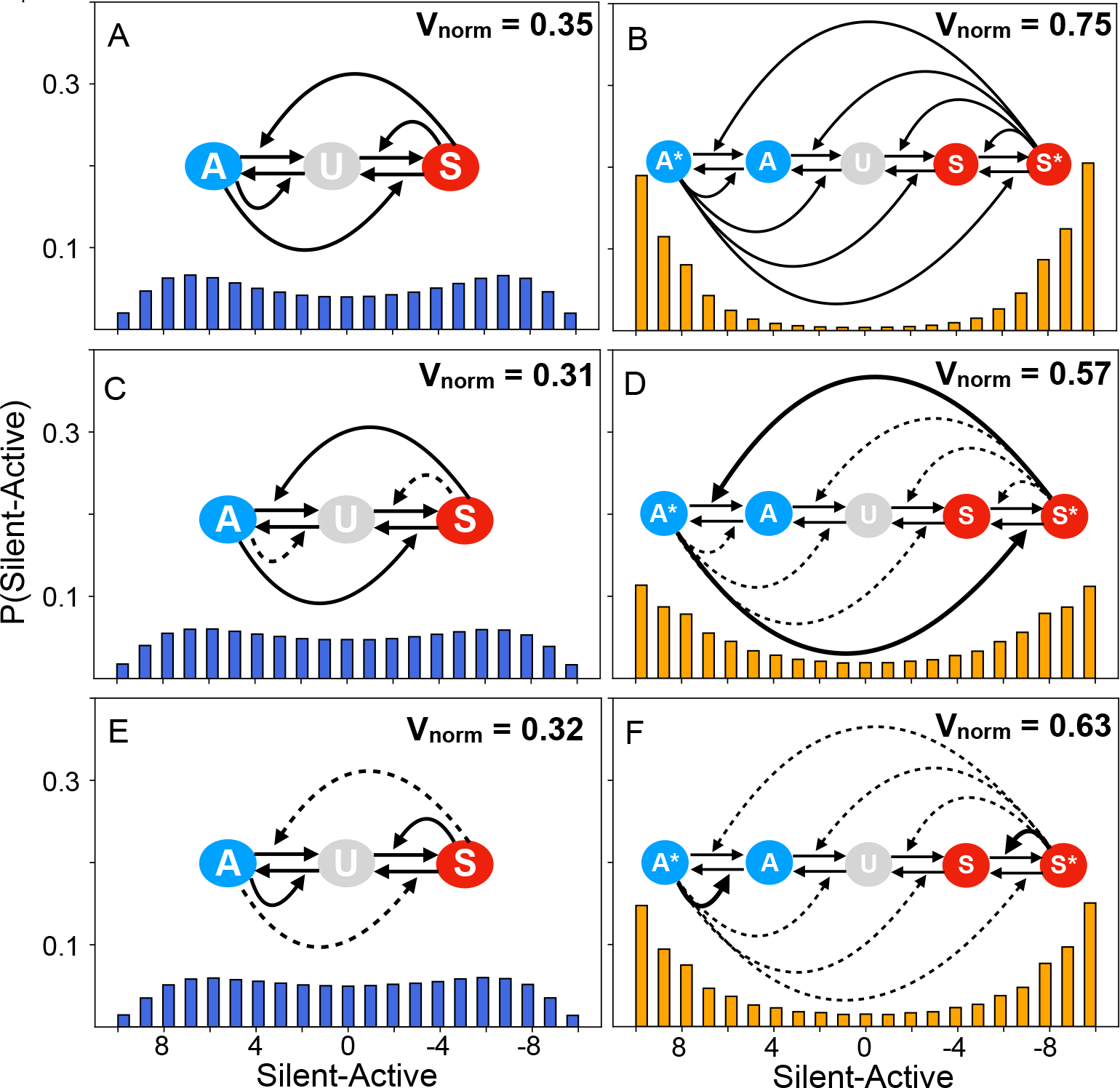
Additional recruitment steps in models with non-local interactions and positive feedback from both sides (A/A* state and S/S* state) increases robust bi-stability. A) shows the behavior of the classical three-state model with only global recruitment (solid arrow). At each update-step, all recruitment types occur with equal probabilities of 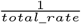 and all spontaneous conversion attempts occur with a probability of 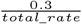 (30% of the rate of recruitment attempts) where *total_rate* = ∑ *X_n_* and *n* being the number of reactions. B, C) As in A but with a combination of local and non-local recruitment. Dashed lines mark reactions that only occur between neighbor nucleosomes. Three versions of three-state models (left panels) result in similar probability distributions with comparable variances (Var) in the number of silent-active nucleosomes. The analogous 5-state models (panels B, D, F) have a much higher value of Var, approaching the maximum bi-stability of Var= 100 for the L=10 nucleosome system. See also Fig. S4.

Due to the increased number of states and reactions of different types (local- and global positive feedback conversions), the subsequent models are simulated by a Gillespie-type update.

We performed the simulation at a relatively high fraction of spontaneous conversion attempts (30%) compared to actual recruitment attempts. The degree of bi-modality is quantified through the normalized variance sampled over a very long simulation

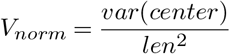

where *len* = 10 is the number ot nucleosomes, the A state is assigned a value of −1, and the silenced state (S*) is assigned a value of +1. In Figure 2, *center* = 10 is also the number of nucleosomes. The term “center” is used because later explorations consider the central subset of a larger system. The normalized variance (*V_norm_*) is shown in the upper left corner of each panel. The higher *V_norm_*, the lower the switching rate between the two-stable states. Generally, all three-state models (Fig. 2 A, C, and E) are marginally bi-modal with a variance *Var* ~ 30-35% of its maximal possible value, while the 5-state models (Fig. 2 B, D, and F) have a substantially higher variance of 57-75% marking robust bi-stability. One also notices weaker bi-stability for models with local feedback reactions that attack states more locally in the recruiting state space (Fig. 2 C, D).

The results demonstrate that additional intermediate states increase the robustness of bi-stability, even if only a small subset of the feedback reactions are non-local (Fig. 2 C-F). Moreover, the probability distributions and variances of 5-state and 7-state models are similar to three-state models with a squared or cubed spontaneous conversion parameter *β* (Fig. S4).

The number of spontaneous reactions converting one nucleosome across *n* — 1 steps involves at least *n* — 1 successful spontaneous conversions towards the minority state (ignoring unlikely recruitment from the minority state). Thereby one effectively re-scales the spontaneous conversion *β* with a factor ∝ *β*^*n*-1^. This re-scaling makes spontaneous transitions away from the minority state increasingly unlikely with larger *n*, which in turn causes the more stable bi-stability seen in Fig. 2 and Fig. S4.

Altogether, we predict that small patches of nucleosomes may realistically exhibit bi-stable and epigenetic memory in themselves, provided that there are several intermediate states subject to non-local positive feedback. And this may be true even if only a small subset of the read-write recruitment reactions reaches beyond the nearest neighbor.

### Combined global negative and positive feedback enables localized epigenetics

Multiple positive feedback loops, including some which facilitate non-local spreading, are necessary for stable maintenance of alternative epigenetic states. However, what prevents a winning state from expanding across the entire genome has remained a puzzle. In the winner takes all dynamics associated with global positive feedback, preventing the winning state from spreading across all boundaries is difficult. Thus, the confinement of epigenetically stable states requires additional mechanisms that prevent unlimited spreading.

Here we address the question of heterochromatic boundaries. For example, in fission yeast, it has been shown that *Epe1*, a putative histone demethylase, is crucial for proper functioning heterochromatic boundaries (16). Interestingly, *Epe1* is recruited by *Swi6*, a reader of the heterochromatic state, and ChIP-seq profiles show that *Epe1* is enriched explicitly in heterochromatin. Moreover, (32) has shown that *Epe1* associates with the *SAGA-complex* that contains several histone acetylases (HATs).

Could models augmented with negative feedback from the silent nucleosome state enable localization of bistable heterochromatic regions? To answer this question, we explored 4-state models with one negative feedback reaction mediated by the S*-state to explore the qualitative behavior of such a mechanism.

After examining models with distance-independent global feedback (see supplement figures S5, S6 and S7) that enable localized bistability but were fragile and sensitive to system size, we used distance-dependent global feedback in simulations. The observation that contact probabilities between nucleosomes decrease as a power law 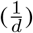 with distance *d* between nucleosomes (33) justifies this choice. The model is based on many known interactions between modified nucleosomes and read-write enzymes (Fig. 3 A) (see supplemental Figure S11 for a description and references). We imagine that a subset of the enzymes could make use of the looping, while others might act only between neighboring nucleosomes.

**Fig. 3.**
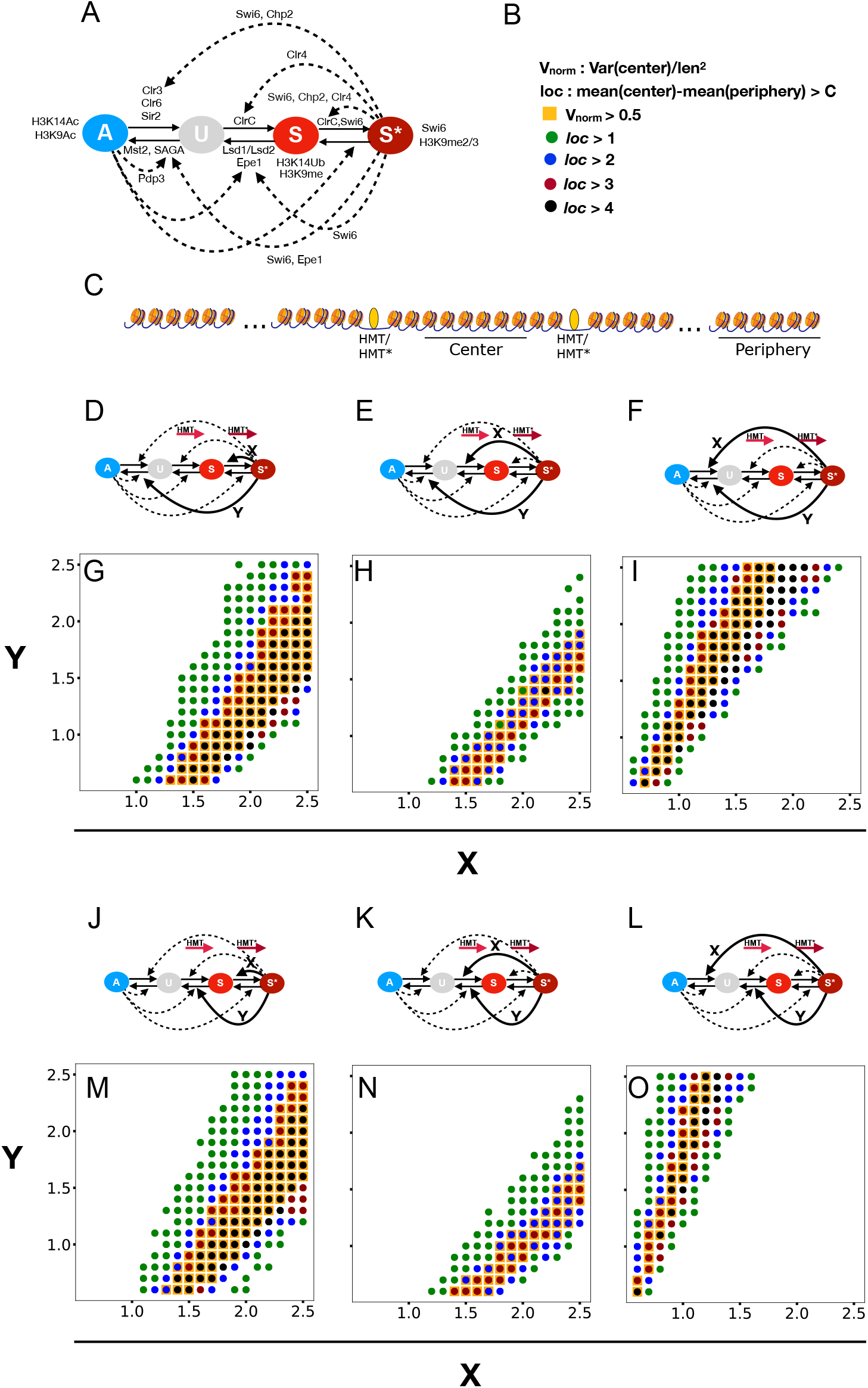
Parameter scanning of models. A) 4-state model structure based on known interactions between read-write enzymes and the modifications they read and write (see supplemental figure S11 for more details and references). B) Bistability criterion and barrier measure. C) Schematic of the system. Simulations considered a 40 nucleosome system without barriers and with two silencers. Fig. 2 D-F) and J-L) illustrate used models. Straight, bold lines mark 1/*d* distance-dependent global feedback, while dashed lines mark local reactions. All local feedback rates are set to a value of 1 and all spontaneous conversion rates to a value of 0.05. The global feedback parameters X and Y were varied in Scans G-I) and M-O). All models can exhibit localized bi-stability. Model G) and M) is most robust as they predicts most working parameters. See also Figures S5 to S8.

The degree of localization of the modification state is quantified by:

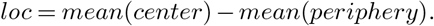

Here, the “center” is the difference between silent and active states for the six central nucleosomes located within the two nucleation sites. The “periphery” is the difference between silent and active states of the six right-most nucleosomes in Fig. 3 C). The “mean” refers to a time average taken over a very long simulation.

The localization measure ranges between 0 (minimum) and 12 (maximum). However, since we are interested in localized bistability, a value of *loc* = 6 is already extremely high. Thus, the *loc* measure has values of 1,2,3 or 4 with larger *loc* indicating more effective confinement of the central region state from the outside regions. Additionally, we demanded that *V_norm_* for the center be larger than 0.5 to count a parameter set as bistable (Fig. 3 B). This approximately corresponds to a distribution like the one seen in Fig. 2D.

We did extensive parameter scanning of several model versions with one distance-dependent global negative- and one distance-dependent global positive feedback, respectively. In each case we tested for parameter values that provide confinement and bistability (Fig. 3 D-O). All tested models show regions of simultaneous confinement and bistability. However, the most robust models, in terms of the largest overlapping parameter space that enables localized bi-stability, were models with S*-mediated distance-dependent global StoS* transitions (Fig. 3 D and J).

Fig. 4 shows simulation results of the same model as in fig. 3 J but with with a system size of L=80 nucleosomes instead of L=40 nucleosomes. This figure illustrates that the model with depicted parameter values (Fig. 4 B) provides balanced prolonged active and silenced states between two silencers for several generations and stochastic switches between both meta-stable states. This results in a relatively high normalized variance *V_norm_* of 0.68 and a mean around zero, as well as a low peripheral *V_norm_* of 0.03 and a low mean of −5.5 (Fig. 4 B). Stochastic switches between stably active and silent states happen because the random appearance of A-nucleosomes within the silencers may amplify themselves through local positive feedback, leading to a switch from silent to an active region state with protection from conversion attempts by the S*-state. On the other hand, when the region between the silencers is in the active state, randomly appearing S*-nucleosomes result in switches from active to silent region states with the S*-mediated StoS* conversions protecting themselves from A-nucleosomes while simultaneously preventing UtoS transitions in the surroundings to ensure localization. Fig S9 shows that we can sustain robust localized bistability despite regular disruptions caused by DNA replication.

**Fig. 4.**
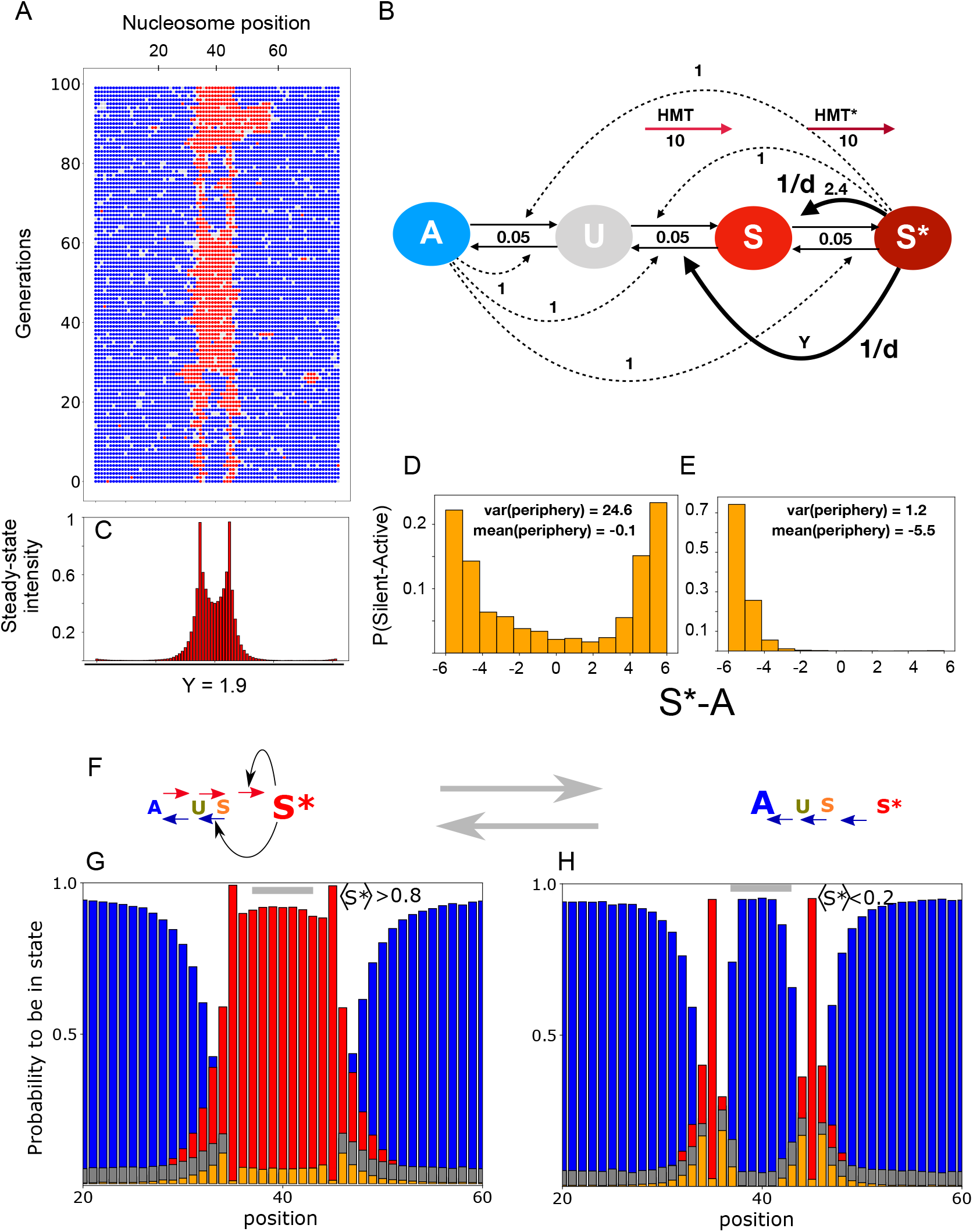
Localization & bistability within two silencers using 1/*d* distance-dependent global negative- and positive feedback shown in panel B). A) Example time-space plot of the model configurated as shown in B) and corresponding steady-state enrichment profiles of S*-nucleosomes C). D) Bi-modal distribution of nucleosome states within the two silencers. E) Distribution of states outside the silencers. F) The active global feedback and dominating local recruitment reactions (← or →) in silenced (left) respective active state (right). G) combined with F) illustrates how S* changes StoU on just outside the silencers where U is present. This in turn allow further recruitment to an A state by a neighboring A (blue). As a result the silenced region represses its own spreading. Panel H (with F) show a dominating A state that maintain itself by local recruitments (blue ←). See also Fig. S9.

Altogether, we find that 4-state models with combined global negative and positive feedback enable localized bistable regions surrounded by euchromatin. Furthermore, this sharp separation between silenced and active regions of the genome did not need explicit barriers but instead required silencers in the form of strongly bound transcription factors.

In (31), transcription has been suggested to act as a global positive feedback from the active side. Thus, the binding and elongation of RNA polymerases restricted to promoters and gene bodies could naturally confine bistable regions. Our analysis here focuses instead on local feedback from active nucleosomes and is not limited to genes but includes other genomic domains like intergenic regions. Our approach does not exclude the transcription option if global positive feedback towards the active side mediated by transcription takes place all over the region encapsulated by the two silencers and is confined to this domain. Thus transcription in the persistently active region surrounding the silencers in Fig. 3 C should not directly influence histone modifications in the region encapsulated by the silencers.

### Global negative feedback is consistent with shape and size of non-centromeric H3K9me-profiles

Since many noncentromeric H3K9me3 domains in mammalian cells and heterochromatic islands in fission yeast have ChIP-seq profiles with localized peaks and soft borders (16), we asked whether our model can recapitulate these features. In Figures 1, S1, S2 and S3 we discuss the classical model for this phenomenon (18), pinpointing its weaknesses and a possible repair in terms of multi-step recruitment processes. Here we suggest a scenario that is also robust to long-range positive feedback around the central inducing recruiting silencer and a relatively high degree of direct conversion attempts (5% of feedback attempts).

Fig. 5 revisits the dynamics of the model from Fig. 4 and demonstrates that localized peaks of modifications can also be supported by increasing the strength of a long range negative feedback. This allows for localized methylation profile even in presence of long range positive feedback and high direct conversion rates (of 5% of recruitment rates). Fig. 5 D-F) illustrate that Spatio-temporal fluctuations are large and persistent in time. Thus through the regulation of nucleosome modifications, we predict substantial cell to cell variations in gene expression (34) for promoters at moderate distance to such nucleation sites, and levels of produced proteins (35). We speculate that indirect gene regulation through epigenetic histone marks may provide yet another path to bursty gene regulation, beyond effects from persistent supercoiling variations (36, 37) or strongly binding transcription factors (35, 38).

**Fig. 5.**
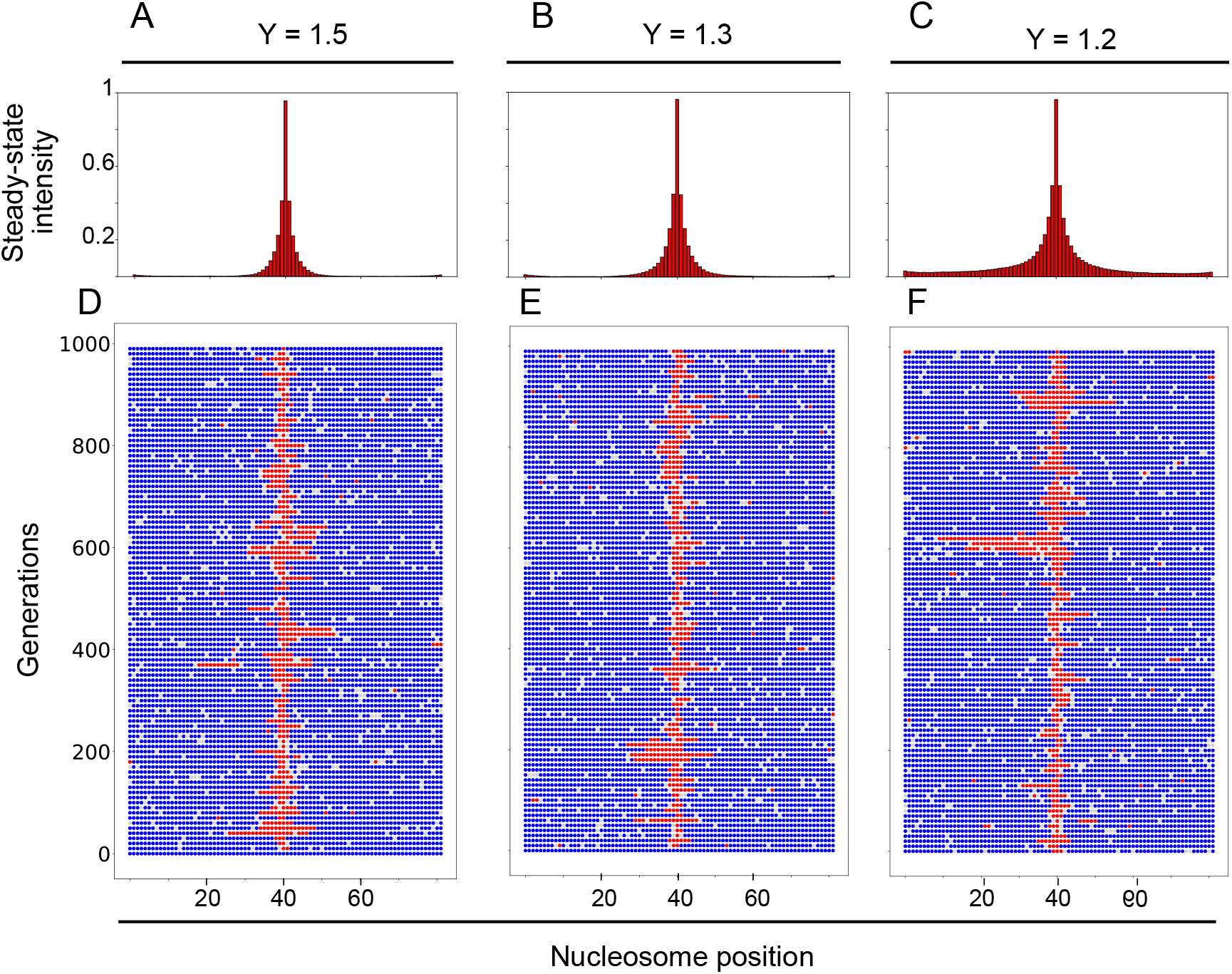
Model from Fig. 4, but applied to a system with only one central nucleating site. One sees that such a system shows confinement when negative global feedback Y is large enough. This illustrates that observed nucleosome state profiles (18) can be obtained even in the presence of substantial non-recruited conversions and long range positive feedback. See also Fig. S10.

### Comparison with experimental data

Consistent with our analysis, Epe1 is recruited by Swi6 and is enriched at heterochromatic domains like the mating-type locus, pericentromeres, and telomers, as well as heterochromatic islands (39) (16). More broadly, our model structure is consistent with known properties of read-write enzymes like positive feedback due to binding to nucleosomes modified with their substrate and subsequent allosteric activation of their catalytic activity leading to modifications of nearby nucleosomes (See Figure S11). Additionally, our modeling is consistent with the typical size of heterochromatic islands (around 3 kb) (39), and might explain why almost all of the *S. Pombe* genome is enriched with acetylated histone modifications (euchromatin) (40, 41).

Another experimental finding that agrees with our analysis is the ectopic spreading of heterochromatin in Epe1 deletion strains (42), as well as the expansion of heterochromatin at heterochromatic islands and the appearance of entirely new heterochromatic islands upon Epe1 deletion (39).

An interesting observation of our modeling is that it can produce occasional ectopic spreading outside the confined region. This might explain the so-called position effect variegation, the spontaneous and stochastic silencing of genes incorporated into genomic loci close to heterochromatic regions (43).

Although bistable chromatin has been observed at the shortened mating-type region in *S Pombe* Δ*K* strains, direct evidence for bistability of small regions in *S Pombe* is still missing. However, direct evidence for bistability of small (2-4 kB) chromosomal regions exists in other organisms like *budding yeast* (8) or *Arabidopsis thaliana* (12). Our model predicts that suitable nucleation sites (around 10-15 nucleosomes) might generate bistable chromatic states in the encapsulated area when positioned close to each other. In the future, it will be interesting to test this hypothesis experimentally in *S Pombe* by, e.g., creating an artificial locus with nucleation sites like the *Atf1* binding site within the mating-type locus or different nucleation sites of heterochromatic islands. Finally, epigenetic inheritance in bistable epigenetic regions often requires silencers or other protein-based memory elements (8, 13, 44) in addition to positive feedback. Interestingly, a recent study in budding yeast by (8) showed that the degree of bistability depends on both the degree of nucleation at silencers and positive feedback via recruited nucleosome conversions. In that study, the degree of nucleation at silencers has been modulated by deletions of several proteins that bind to the silencers E and I of the *HMRα* locus and which recruit the SIR-complex (fig. 6 B). The SIR complex consists of three subunits, SIR3, SIR4, and the histone-deacetylase SIR2. In budding yeast, the heterochromatic state (S*) is characterized by a deacetylated and SIR-bound nucleosome state. Specifically, silencing occurs due to recruitment of the SIR complex via nucleosomes that don’t carry any active modifications (specifically H3K79me and H4K16ac) and subsequent deacetylation of nearby nucleosomes (45) (see Fig. S12 B,C).

**Fig. 6.**
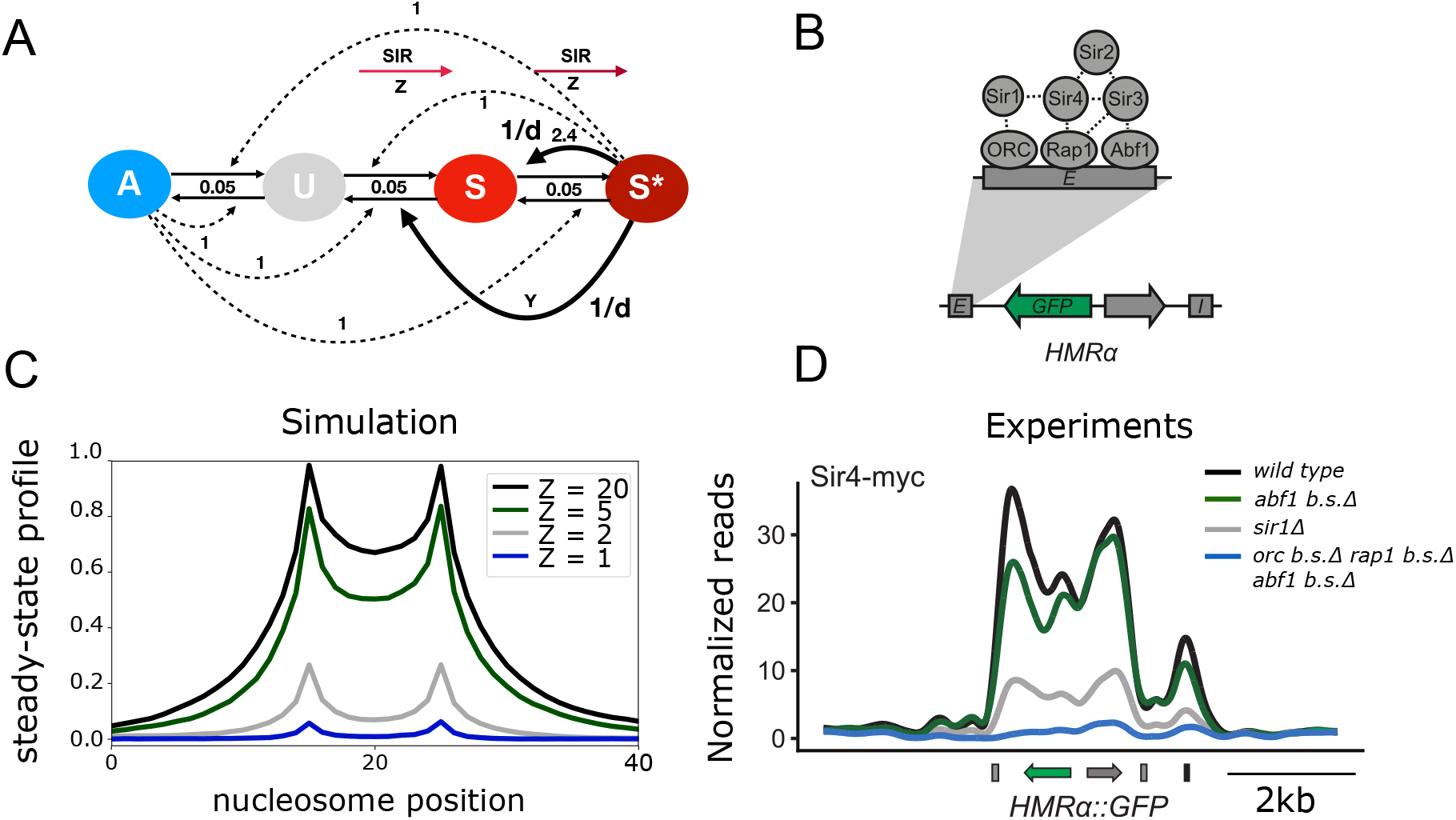
Comparison of steady-state profiles and experimental ChIP-seq reads in *S. Cerevisiae*. A) Same model as in Fig. 4 and Fig. 5 but with fixed S* mediated negative feedback and variable recruitment strength at the silencers (*SIR* = *Z*). Nucleation happens via local recruitment of the SIR complex to the silencers (see text and Fig. S12). B) Schematics adopted from (8) showing the different proteins involved in the recruitment of the SIR complex to the silencers. C) Steady-state profile of time-averaged S*-nucleosome enrichment from simulations of the model and parameter values shown in A and with different degrees of direct SIR recruitment (SIR) at both silencers. Panel D is adapted from (8), showing ChIP-Seq profiles of Sir4-myc enrichment on the *HMRα::GFP* locus of different *S. Cerevisiae* mutant strains where different combinations of boundary elements have been mutated. See Fig. S12 for more details

We tested our model against the findings of the above results (8). To do this, we first identified parameters that predicted the whole region within both silencers to be almost entirely silenced while leaving the region outside the silencers active (Fig. 6 A). Subsequently, we weakened the nucleation strength (see Fig. 6 and Fig. S12). In agreement with the experimental results of (8) (Fig. 6 D), we found that the nucleation strength indeed determines the steady-state profiles of SIR4-bound nucleosome enrichment and the degree of bistability within the silencers (Fig. 6 C and Fig. S12).

Interestingly, mechanistic crosstalk between different types of active modifications and their read-write enzymes has been recently elucidated (46, 47). This crosstalk suggests multistep positive feedback processes from active and silent nucleosomes (see Fig. S12 B). However, negative feedback remains to be shown in budding yeast. Our analysis suggests that one should search for additional read-write enzymes in this organism. Notice also that there are sharper boundaries between SIR4-enriched nucleosomes in experiments compared to simulations (Fig. 6 C, D). A simple extension of our model with an additional nucleation of active modifications close to the silencers but outside the *HMRα* region enables sharper boundaries in simulation due to synergy with the nucleation of silent nucleosomes at the silencers (see Fig. S12). This is in line with the observation that active promoters close to the E and I silencers help prevent the ectopic spreading of Sir4 (48) to the surrounding euchromatic regions.

Overall, our theoretical analysis suggest that negative feedback could play a role in creating heterochromatic boundaries at the *HMRα* locus. It will be interesting to see whether such negative feedback exists as a part of a mechanism to prevent ectopic propagation of the SIR complex or whether boundaries in this organism rely on different mechanisms.

## Discussion

Localization phenomena are found across the natural sciences, from geology to physics and to life. In biology, it is associated with the specific control of/competition between the many diverse life processes. Within gene regulation, one may well argue that epigenetics in trans works so well by diffusive recruitment of regulators to promoters (49), that there is no need at all to bother about the more localized epigenetics in *cis* that may be provided through nucleosomes (50). However, specificity and the ability to store the memory of cellular states locally on the genome will increase the computational power within a cell. For example, it allows the observed differentiation into hundreds of different olfactoric neuronal cells by using local positive feedback provided by nucleosomes read-write enzymes (51, 52). Furthermore, the spreading of histone marks from DNA-bound transcription factors may well facilitate the regulation of promoters, thus combining the best of the *cis* and the *trans* mode of regulation (28, 53, 54).

The current paper explored the ability to localize epigenetic marks around an inducing factor, for example, a silencer or a transcription factor. Across the genome, a main effect of nucleosome modifications is to maintain an externally regulated state in a 5-20 nucleosome region surrounding typical promoters. See ref. (55) for a review that points out that histone modifications can be either cause or consequence of genome function depending on the context. A simple setup of this type was provided by the one-step model (Fig. 1A) where one modification type is allowed to propagate between neighbors by locally acting read-write enzymes (18, 19) (Figures 1, S1 and S2). With the additional assumption of the complete absence of non-recruited conversions like direct methylation, the model reproduces the observed localization of modifications with a spatial extension given by a ratio of spreading rate to decay rate for the modification in question. However, in a more realistic setting, each nucleosome is expected to be also modified by enzymes that are not only associated with neighbor nucleosomes. These non-local interactions break the confinement and lead to varying degrees of unbounded escape from the region of interest (see Figures 1, S1 and S2 D).

One obtains a more robust local spreading when one considers that nucleosomes in opposing epigenetic states are separated by more than one enzymatic reaction in the nucleosome state-space. Just adding one intermediate state, one finds a finite threshold against direct conversions (see Figures 1, S1 and S2 G, H). Fig. 5 demonstrates that possible negative feedback allows for localization even in the presence of long range positive feedbacks. Importantly, then multistep separation of silenced and active nucleosome states with long-range positive feedback also opens for robust bistability (3), detailed further in Figs. 2–4.

Metastable epigenetic states of systems of nucleosomes are seen in many real-world situations, from mating-type regions in yeast (1, 2), over the memory of winter (vernalization) in plants (12) to the multiple states of olfactoric neurons in mammals (51). In these cases, the chromosomal region of concern is limited, and the winning epigenetic state is prevented from spreading across the chromosome. Such localization is difficult to obtain with only positive feedback, as it typically favors a run-away effect along the genome. This run-away effect is especially the case if we accept that some of the recruitment processes have to act non-locally along the genome (3), and thereby typically will tend to bypass any localized barriers.

Combining positive and negative feedback is common in biology. In confinement and associated pattern formation, it has been suggested that Turing mechanisms play a role in the development of some organs (56, 57). Also, one has a Turing-like combination of local acting positive with a more global negative feedback (51) acting through AdCy3, and LSD1 downregulation (52). This mechanism allows the cell to select one and only one expressed olfactoric receptor protein.

Here we considered combined feedback of read-write enzymes between some fixed inducers or barriers on the chromosome. We found that such feedback could provide localized bi-stable regions, provided that the negative feedback was long-range. Noticeably, there is a candidate for negative feedback using *Epe1* along the H3K9 modification axis in *S. pombe*. Currently, there is literature support for both indirect acetylase activity (as in Figs. 3 D-F, S5 H and J, S6 and S8 bottom) and a direct demethylase activity (as in Fig. 3 J-L, figs. 4–6 and fig. S5 D and fig. S8 top). Our analysis makes both scenarios workable.

A sampling of model variations suggests a need for long-range enzymatic conversions for both positive and negative feedback. However, there is also a strict need for some local positive feedback (4–6). We found that localized bi-stability was easier to obtain when most positive feedback reactions were more localized than negative feedback reactions (see Fig. 4F, G).

Interestingly, recent evidence suggests that liquid-liquid droplet formation (phase separation) via Swi6^HP1^ might contribute to the formation of heterochromatin under certain conditions (58, 59). We speculate that phase separation might be a way to render a subset of Swi6^HP1^ target regions bistable by increasing the frequency of recruited nucleosome conversions between linearly distant nucleosomes (60). Moreover, phase separation might even contribute to the confinement of heterochromatic regions, potentially in synergy with nonlocal negative feedback, as suggested in this paper. However, whether phase separation contributes to heterochromatin formation and/or maintenance is still controversial (61), and many questions remain. E.g., how the droplets would maintain a specific size needed for long-term hetero-chromatic silencing is a puzzle. Combining more quantitative experiments of possibly inducible phase separation with polymer modeling could be a way to shed more light on a potential mechanism.

## Conclusion

Overall, our theoretical findings can be summarized as follows:

- Nonprocessive recruitment of read-write enzymes via eu- and hetero-chromatic nucleosomes and a subsequent conversion of neighboring nucleosome states (multi-step positive feedback) allows for increasingly robust confinement in linear spreading models.
- Multi-step positive feedback increases the bi-stability and robustness of models, and this is even the case for models that only have one non-local feedback.
- Long-range negative feedback enables further confinement.
- Distance dependence on non-local recruitment processes significantly increases the robustness of confinement.
- Multi-step processes agree with observations of modification profiles and allow for controlled bi-stability through changed nucleation strength.

We particular highlight the new suggestion that localized silencing or bi-stability could be obtained by supplementing positive feedback with a single negative feedback. A Negative feedback with relatively strong activity directed from the inside to the outside of the silenced region. Candidates for such enzyme activities could be Epe1 or Mst2.

## Methods

### Basic algorithm of purely local two-state models (Figures 1 B, S1 and S2)

To ensure comparability with the model introduced in (18), we used a simple stochastic algorithm (SSA) to simulate the models presented in figures 1, S1 and S2. We simulated the model on a one-dimensional lattice with 257 sites, each representing a nucleosome that can be in one of two states, H3K9un or H3K9me. Simulations were performed for 50.000 update steps, starting from all nucleosomes in the H3K9un state, to ensure that the system has reached steady state.

The steady-state profiles show the time-averaged proportion of H3K9me nucleosomes at each lattice site. A nucleation reaction, a propagation reaction, a demethylation reaction (H3K9me to H3K9un) and a direct methylation reaction (H3K9un to H3K9me) is executed at each update step. Both nucleation and propagation reactions happen with a rate of *k+*. Demethylation reactions are fixed at a rate of *k*— = 0.05 and direct methylation reactions are fixed at *k + direct* = 0.001.

For the nucleation reaction, only the central nucleosome is attempted to be methylated, whereas the propagation reaction attempts to methylate every nucleosome that is the nearest neighbor of an already methylated nucleosome at each update step. A turnover reaction (direct methylation or demethylation) consists of a specific direct conversion of each nucleosome in the system with the above-specified rate.

### Basic algorithm of purely local three-state models (Figures 1 C, S1 and S2)

The three-state model version (Fig. 1 C) is simulated similar to the two-state model but with an additional H3Kac state, so that each nucleosome now exists in one of three states at each step. Simulations were performed for 100.000 update steps starting from all nucleosomes being in the H3Kac state. Also there are now double as many different reactions. A nucleation reaction, two propagation reaction, and four direct modification attempts (an acetylation (H3K9un to H3Kac) and a deacetylation reaction (H3Kac to H3K9un) in addition to the demethylation and direct methylation reaction of the two-state model) are executed at each update step. Both nucleation and propagation reactions happen with a rate of *k+*. A nucleation reaction consists of an attempt to directly convert the central nucleosome to the H3K9me state. For the feedback acting on H3K9un nucleosomes, all H3K9un nucleosomes that are a nearest neighbor of an H3K9me nucleosome are attempted to be converted to an H3K9me nucleosome. In case of a feedback acting on H3Kac nucleosomes, all H3Kac nucleosomes neighboring H3K9me nucleosomes are identified followed by a conversion attempt to an H3K9un nucleosome. Direct modifications are either fixed at a rate of *k*— = 0.05 if the direct transition is towards an active state (H3K9me to K3K9un or K3K9un to H3Kac) or at *k + direct* = 0.001 if the direct transition is towards a silent state (direct deacetylation or direct methylation). A turnover reaction consists of a specific direct conversion of each nucleosome in the system with the above-specified rate.

### Basic algorithm

For all simulations that are not purely local as described above, the system is simulated as an event-driven Gillespie algorithm. Each event is one of *n* reaction types 1,2....*n*, and each event type is either a spontaneous conversion, a recruited conversion, or a nucleation attempt.

First, all rate constants are stored in a list (rates = [*X*_1_, *X*_2_,..., *X_n_*]), and a vector containing the cumulative sums of the rate constants ordered as in the rates-list is created (*cum_rates_* = [*X*_1_, *X*_1_ + *X*_2_,..., *X*_1_ + *X*_2_ +... + *X_n_*]). Then, for each update step, a random number (*rand*) uniformly distributed between 0 and the sum of all rate constants 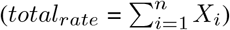 is generated. The first index *k* where *cum_rates_* (*k*) is larger than *rand* · *total_rate_* determines the event that is executed. The absolute time is increased by generating another random number (*rand*_1_) and calculating 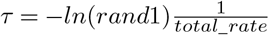.

If a spontaneous conversion is attempted, a random nucleosome in the system is chosen, and a state-change is attempted accordingly (e.g., if a UtoS conversion is attempted, a state change from U to S only happens if a U is selected).

In case of a recruited-conversion, two nucleosomes (*nuc*1 = recruiting nucleosome and *nuc*2 = substrate nucleosome) are chosen and a state-change happens if *nuc*1 and *nuc*2 are compatible with the chosen recruited reaction then conversion is attempted. For example if an S-nucleosome mediated A to U transition (reaction S(AtoU)) is chosen then *nuc2* changes its state from A to U if and only if only *nuc*1 is in the S state and *nuc2* is in the A state).

Nucleation attempts are similar to spontaneous conversion, with the exception that a successful attempt requires the correct position in addition to the nucleosome being in the correct state (e.g., if the nucleation site is at position 40 and a UtoS nucleation is attempted, a state-change from U to S only happens if nucleosome at position 40 is chosen and if it is in the U state).

### Local and global recruitment

In case the chosen event is a local recruitment, *nuc*1 is chosen randomly and a neighbouring nucleosome *nuc*2 is chosen as the left or right neighbour with equal probability. For a distance-independent global reaction, *nuc*1 and *nuc2* are chosen randomly with *nuc*1 ╪ *nuc*2. A distance-dependent global reaction is executed by choosing *nuc*1 randomly and *nuc*2 at distance *d* = |*pos*(*nuc*1) — *pos*(*nuc*2)| with probability 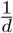 (to the left or to the right of nuc1 with equal probability). A successful move requires *nuc*2 and *nuc*1 to be inside the region. When *d* is chosen, such that *nuc*2 is selected to be outside the region, nothing happens. To secure convergence, we select distance *d* within an interval that is at max the length of the simulated region.

### Barriers

A Barrier is implemented as a nucleosome in a permanent state (T state) that is not receptive to any involved modifications. An example could be that this position on the genome is bound by a specific transcription factor-like, e.g., CTCF.

### Generations and DNA-replication

We defined a generation to consist of 20 conversion attempts of each reaction type (multiplied by the rate constant) per nucleosome on average. We investigated the effect of DNA replication on localized bistability by simulating a state-change of each nucleosome in the system to a U-state with 50 % probability after each generation (Figure S6).

### Analysis

We generated the Probability distributions in figures 2 and S3 from long simulations (10^9^ update steps) starting from all nucleosomes in The U-state. After each update step, the number of silent (S and S*) and active (A and A*) are counted and used for calculating the time-averaged probability distribution of silent-active nucleosomes.

Figures 4 and S6 show histograms of a bigger system subset of 6 respective nucleosomes. We simulated the system for 20.000 generations (about 10^9^ update steps) and recorded the state of each nucleosome after each generation to generate these time-averaged distributions of states.

In the parameter-space plots (Figures 3, S5, S6 and S8) two parameters are varied while fixing the remaining ones. For each parameter set, we run the simulations of 40 nucleosome systems for 10^8^ update steps and test the system’s properties.

First, we highlight situations where the system exhibits bistability (orange); second, we grade the extent to which the system’s interior differs from its exterior. Here interior is defined within the silencing factors, while nucleosomes outside the barrier elements define the exterior.

Occasionally we illustrate the dynamics as space-time plots, displaying all nucleosomes in the system at several discrete time points. These plots allow us to present situations with qualitative differences between the dynamics in regions of the examined system. The time span we show is typically quite long, e.g., in Fig. 5 we show a time equivalent to 1000 generations.

## ACKNOWLEDGEMENTS

We thank the European Union’s Horizon 2020 Research and Innovation Program under Marie Skłodowska-Curie Grant 813282 to J.F.N for financial support. We also thank members of our group and especially our experimental collaborators, Genevieve Thon and her group members, for insightful discussions and guidance.

## Supplementary Materials

**Fig. S1.**
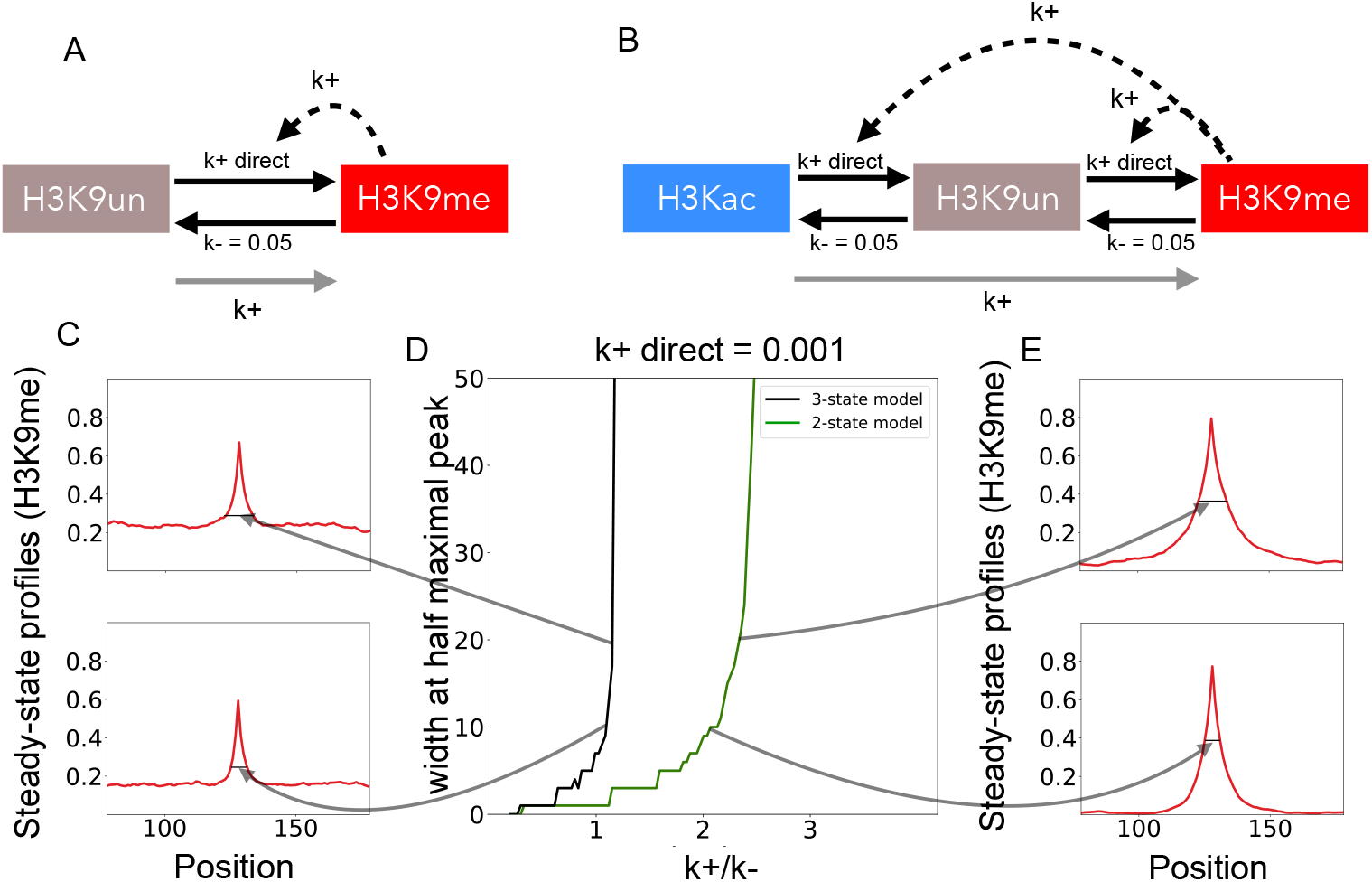
A) An additional recruitment step from the H3K9me-state increases the width of inherently bounded steady-state profiles in the presence of low direct silencing rates. Schematic of a two-state model with local positive feedback from H3K9me-nucleosomes. B) Schematic of a three-state model with local positive feedback with local positive feedback from S-nucleosomes. C) and E) Example steady-state profiles showing the time-averaged enrichment of H3K9me-modified resulting from simulations corresponding to the 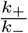 values depicted by the curved arrows. D) Relationship of the width at half maximal peak size as a function of 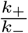 for the two-state and 3-state model with a constant *k*_+_. direct rate of 0.001

**Fig. S2.**
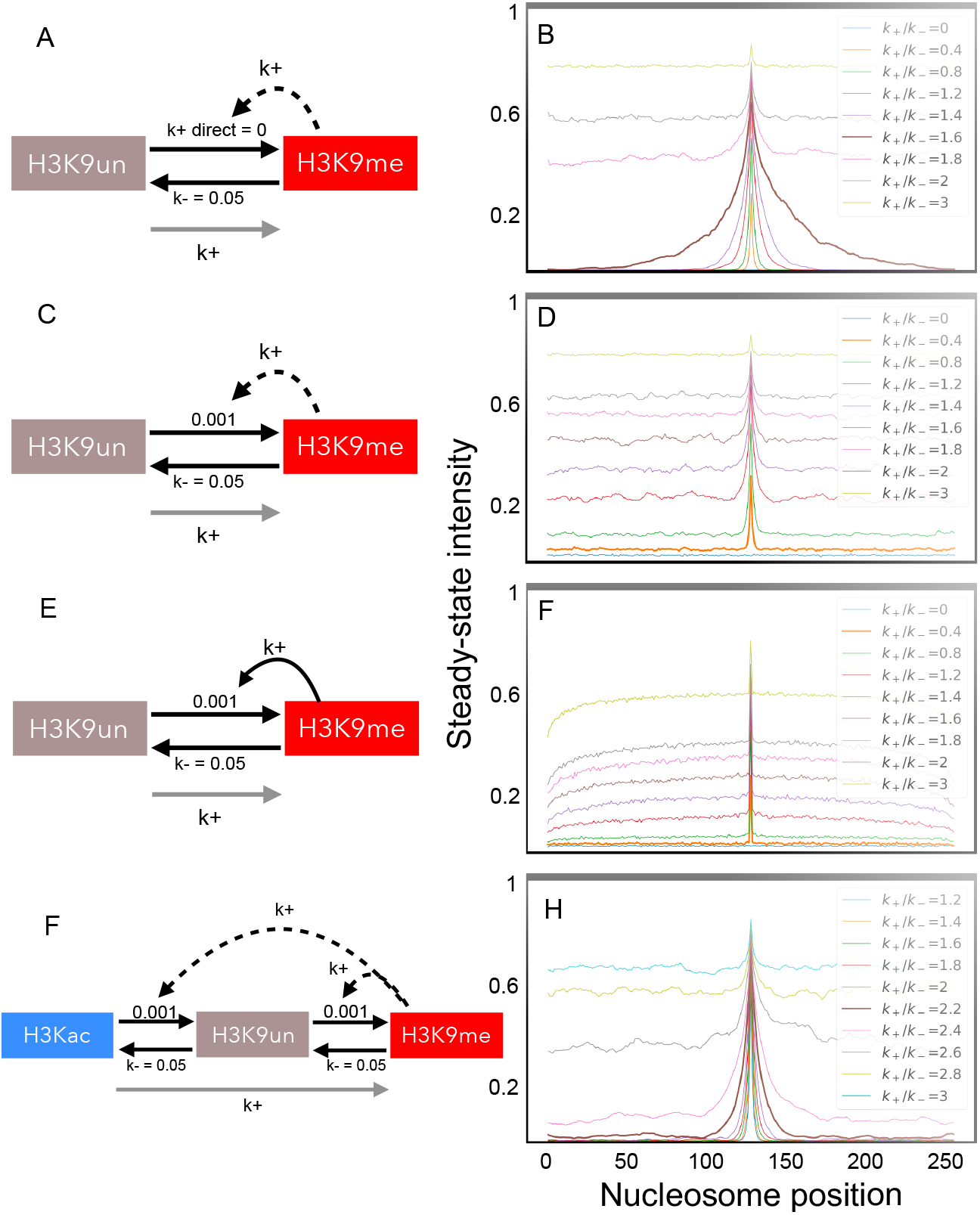
Localization of methylation around a silencer of models with local-positive feedback only from the silenced state. The listed numbers for lines in the figures refer to the *k_+_/k_-_* ratio. A-B) A simple one step model can localize methylation around a silencer in the absence of direct methylation (H3K9un to H3K9me). C,D) The motif is however very fragile to rare direct methylation (that here is 50 times slower than the opposite direction). E,F) Explore a 1/*d* distance dependant recruitment, demonstrating that this does not help on the confinement. G,H) Adding an additional intermediate state increases the robustness towards rare direct methylation, and leads to near complete localization when the ratio *k_+_/k_-_* is below the threshold 2.2.

**Fig. S3.**
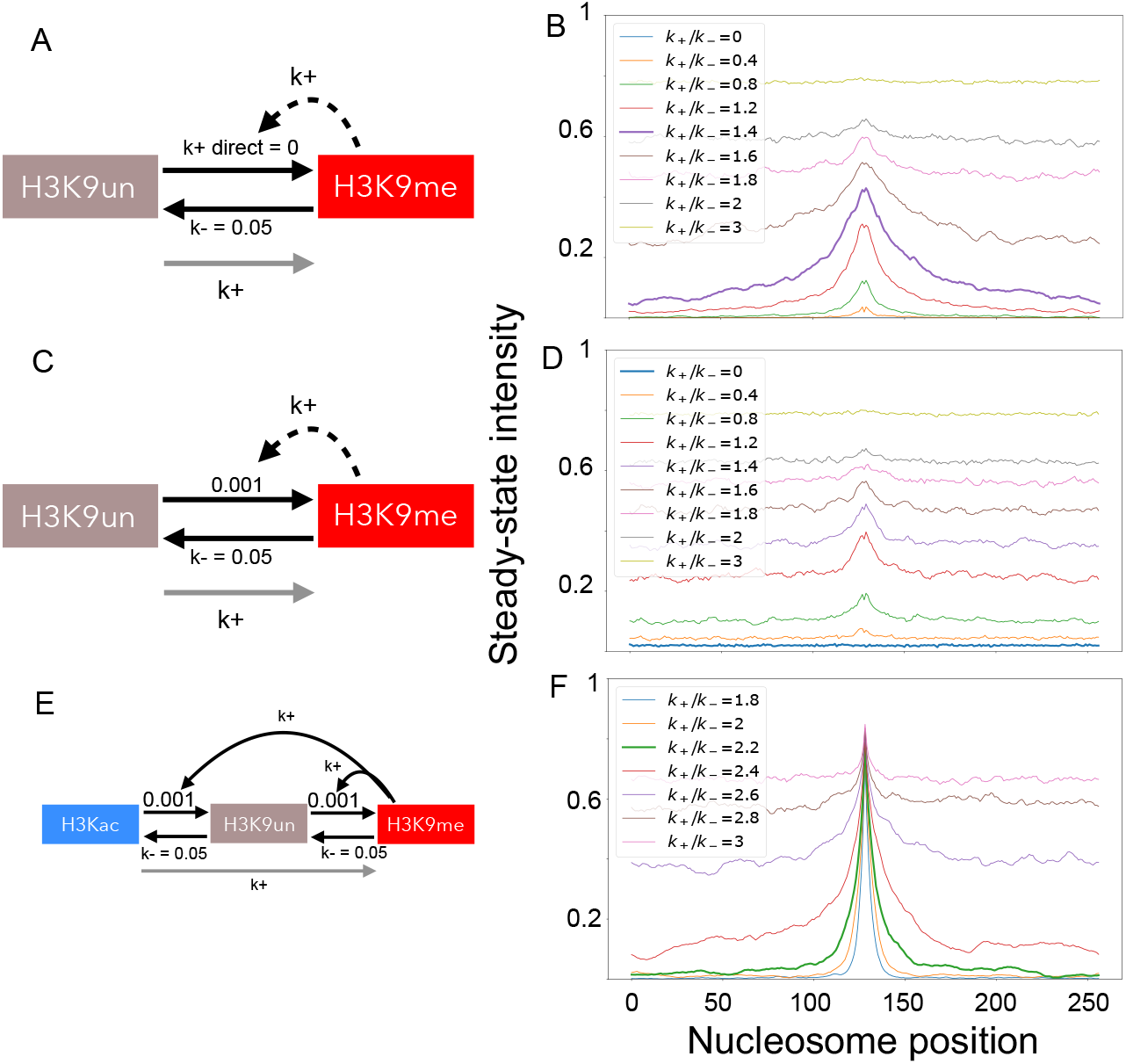
Localization of methylation around a silencer of models with local-positive feedback only from the silenced state. In this figure, spreading from the nucleation site occurs in a distance dependent manner, where nucleosomes are methylated with a probability of 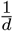 where d is the distance from the central nucleosome. The listed numbers for lines in the figures refer to the *k+/k_-_* ratio. A-B) A simple one step model can localize methylation around a silencer in the absence of direct methylation reactions, similar to Fig. S2 B. C,D) The motif is however very fragile to rare direct methylation of nucleosomes (that here is 50 times slower than the opposite direction). E,F) Explore a three-state model, which similar to Fig. S2 enables confinement below a *k_+_/k_-_* ratio of 1.4.

**Fig. S4.**
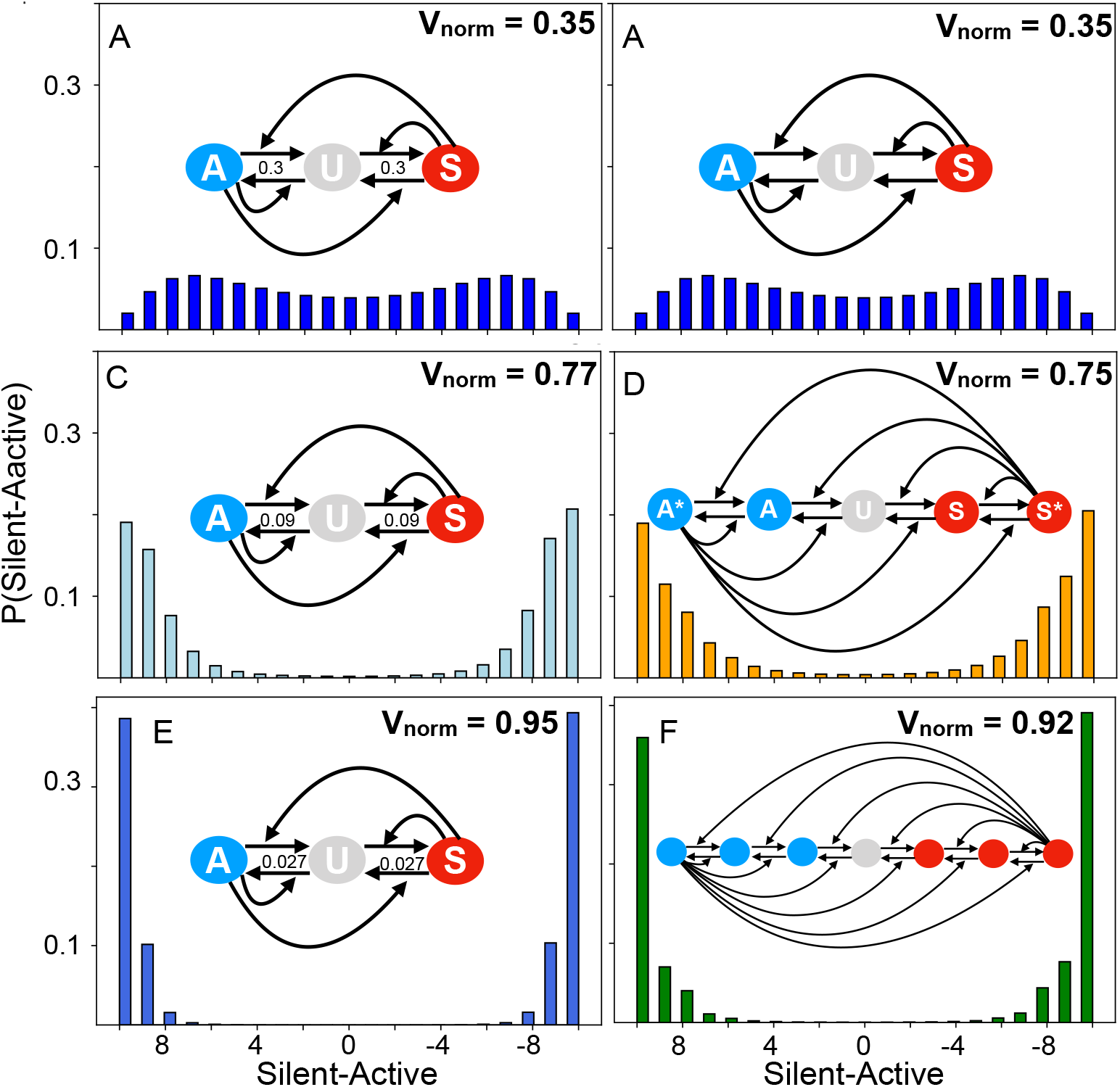
Squared/cubed direct transition rates increase robust bistability similarly as in models with two/four additional nucleosome states subject to positive feedback. The left panels show Silent-Active probability distributions from simulations of 3-state models with a direct conversion attempt-rate constant of 0.3 (A), 0.3^2^ (B) and 0.3^3^ (C). The right panels show Silent-Active probability distributions of a 3-state-model (B), a 5-state model (D) and a 7-state model (E) all with a direct conversion attempt-rate constant of 0.3 and a recruited attempt rate of 1.

**Fig. S5.**
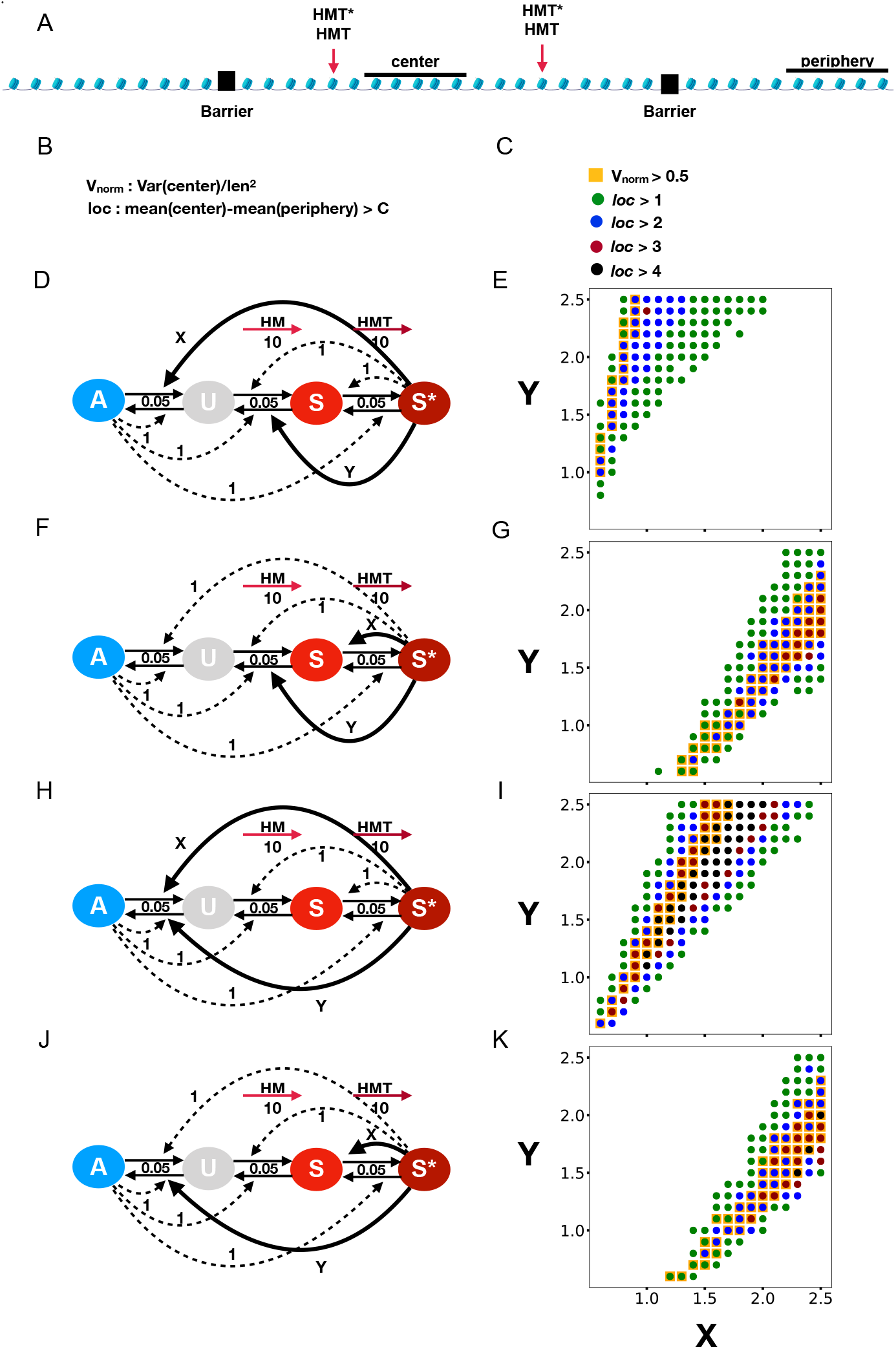
Exploring motifs that provide localized bi-stability. A) schematics of the system comprising 40 nucleosomes, two local barriers and two silencers. In all cases X and Y reactions are assumed distance independent while all dotted recruitment are purely acting between neighbor nucleosomes. One observes that motif H) provides the most robust features (panel I).

**Fig. S6.**
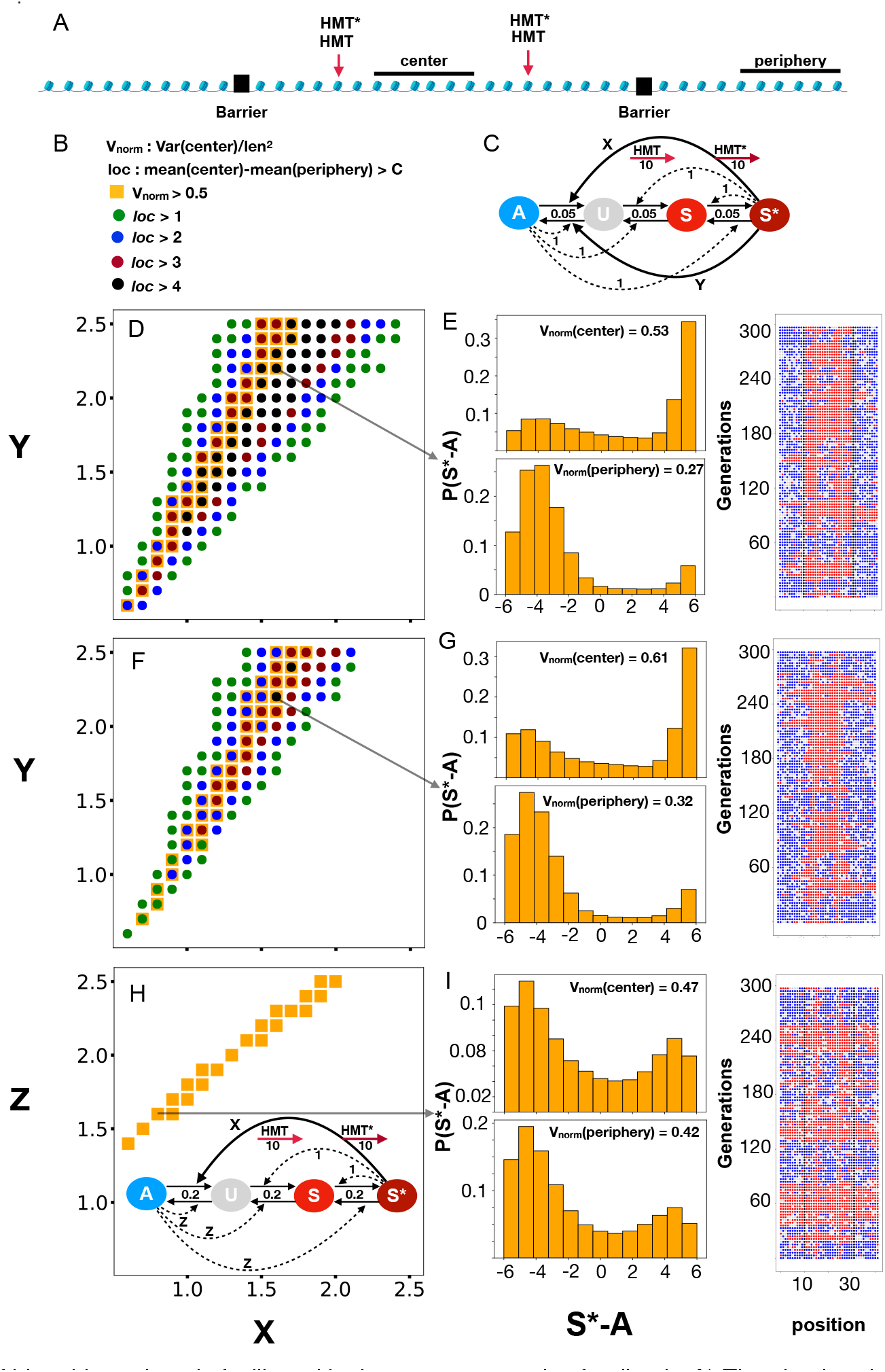
Confinement of bi-stable regions is facilitated by long-range negative feedback. A) The simulated system of 40 nucleosomes, silencers, and potential barriers. B) The interaction rules with solid lines marking non-local reactions and dashed lines depicting local recruitment. D, E) illustrate that localized (colored dots) bi-stability (orange squares) is obtained along with a range of parameters with the strength of negative feedback *Y* proportional to the strength of positive feedback *X*. F, G) Illustrate that such localized bi-stability is also possible when we remove the barriers. H, I) show that localization is not possible in the absence of negative feedback *Y*.

**Fig. S7.**
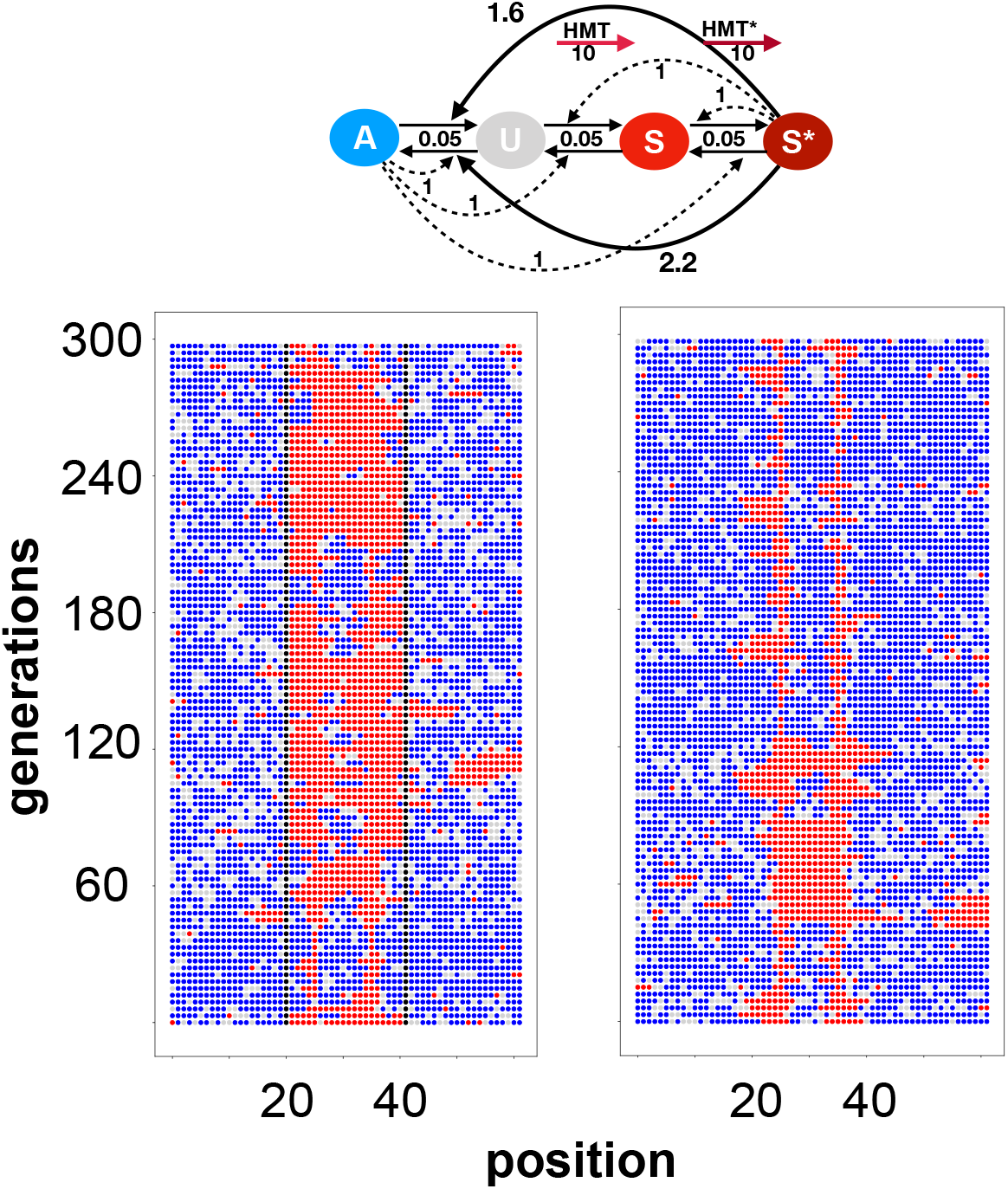
Same model and parameters as in Fig. S6 but on a larger system of 60 nucleosomes. Left panel shows simulation with barriers and right panel shows simulations without barriers. Overall confined bistability is still possible to obtain in larger systems, but confinement is more fragile.

**Fig. S8.**
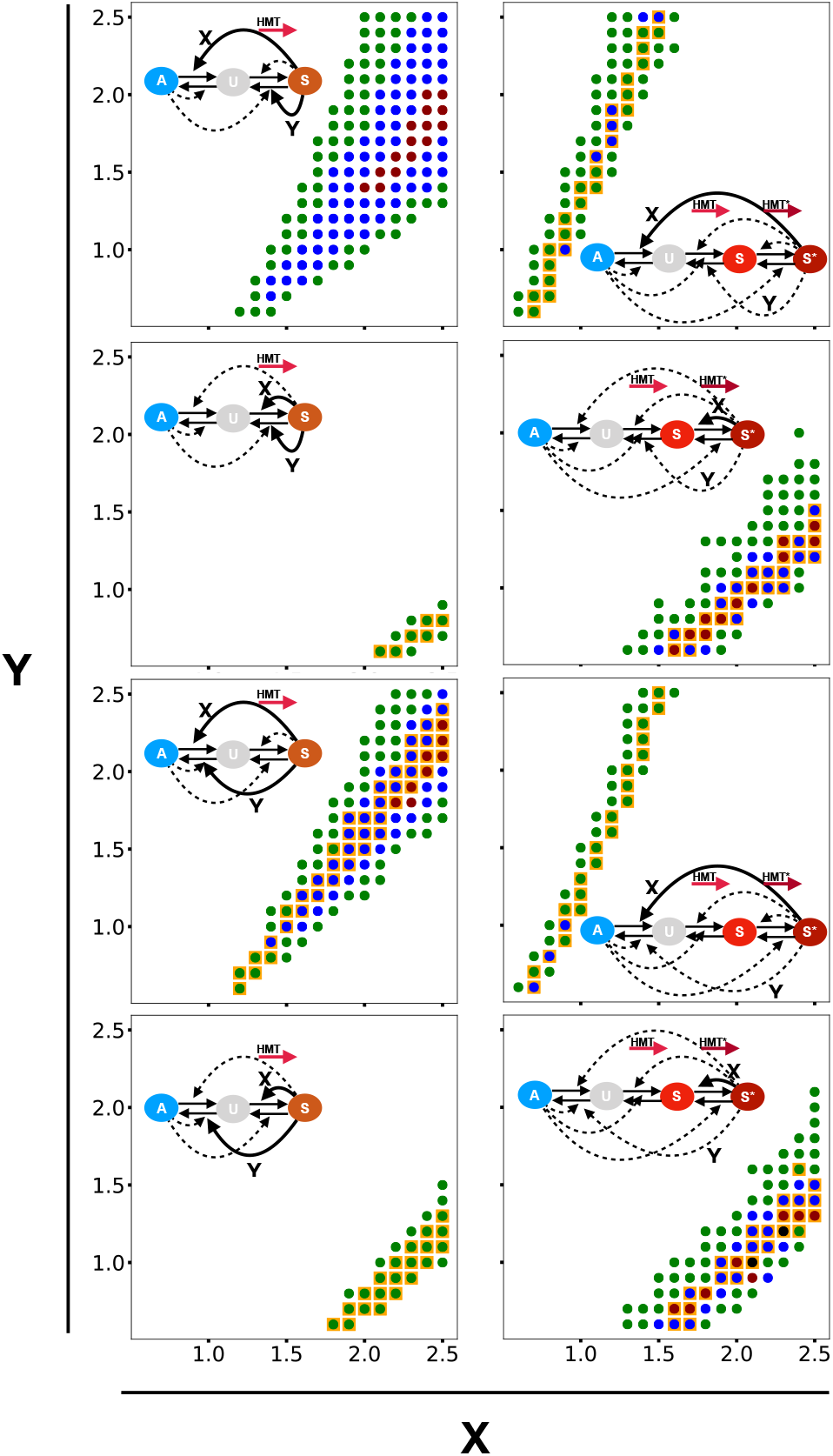
Exploration of alternative motifs with distance dependent global feedbacks. Left panels show that its quite difficult to obtain localized bi-stability between silencers if one only have 3 states. Right panels explore 4 state models where the negative feedback only acts locally. Comparing with Fig. 3 one observes quite marginal confinement and bi-stability.

**Fig. S9.**
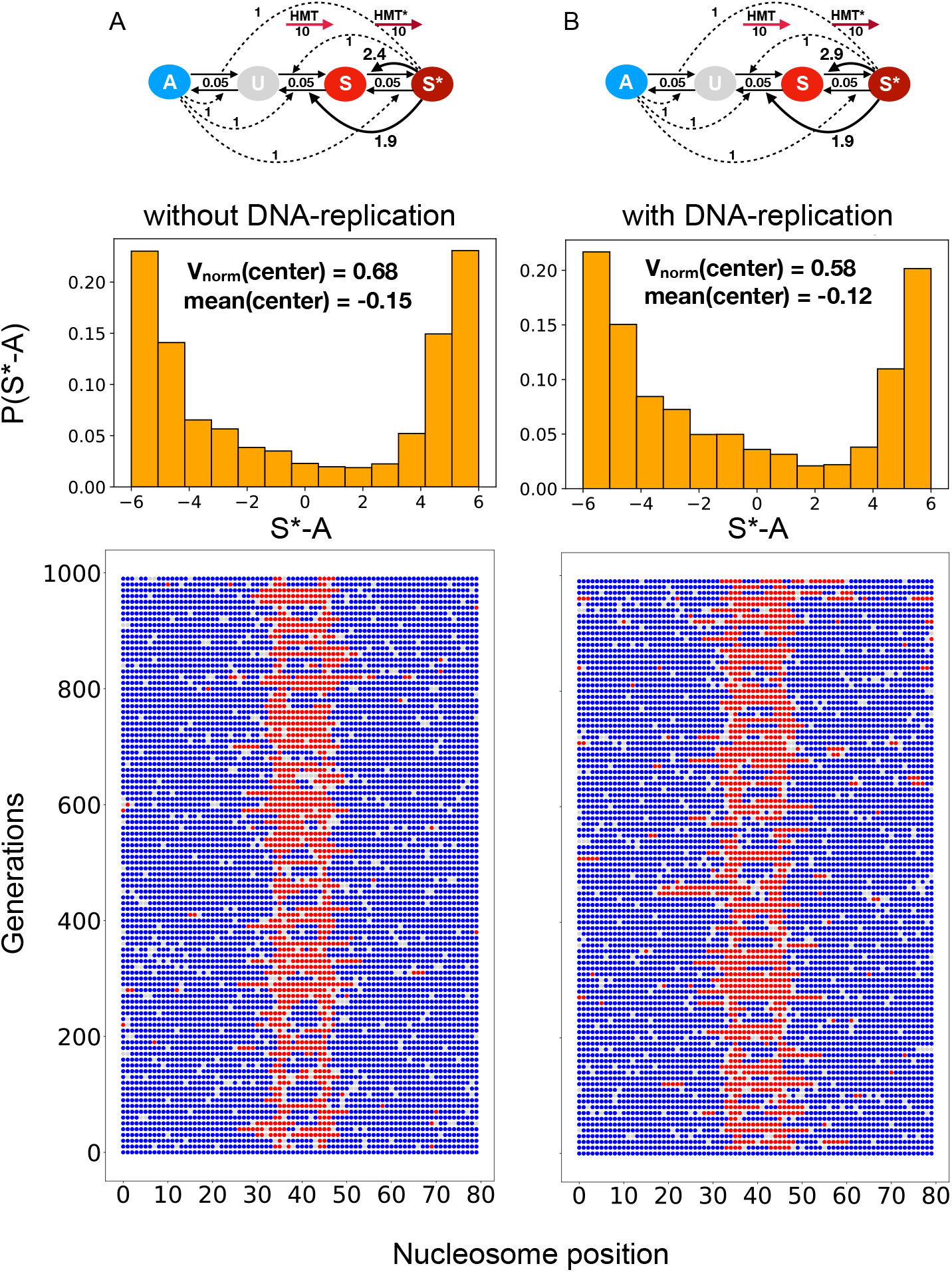
Localized bistability can be maintained despite DNA replication. In the presence of surrounding silencers (nucleosomes with a high nucleation rate), inner localized bi-stable chromatic regions can be maintained despite simulating DNA replication.

**Fig. S10.**
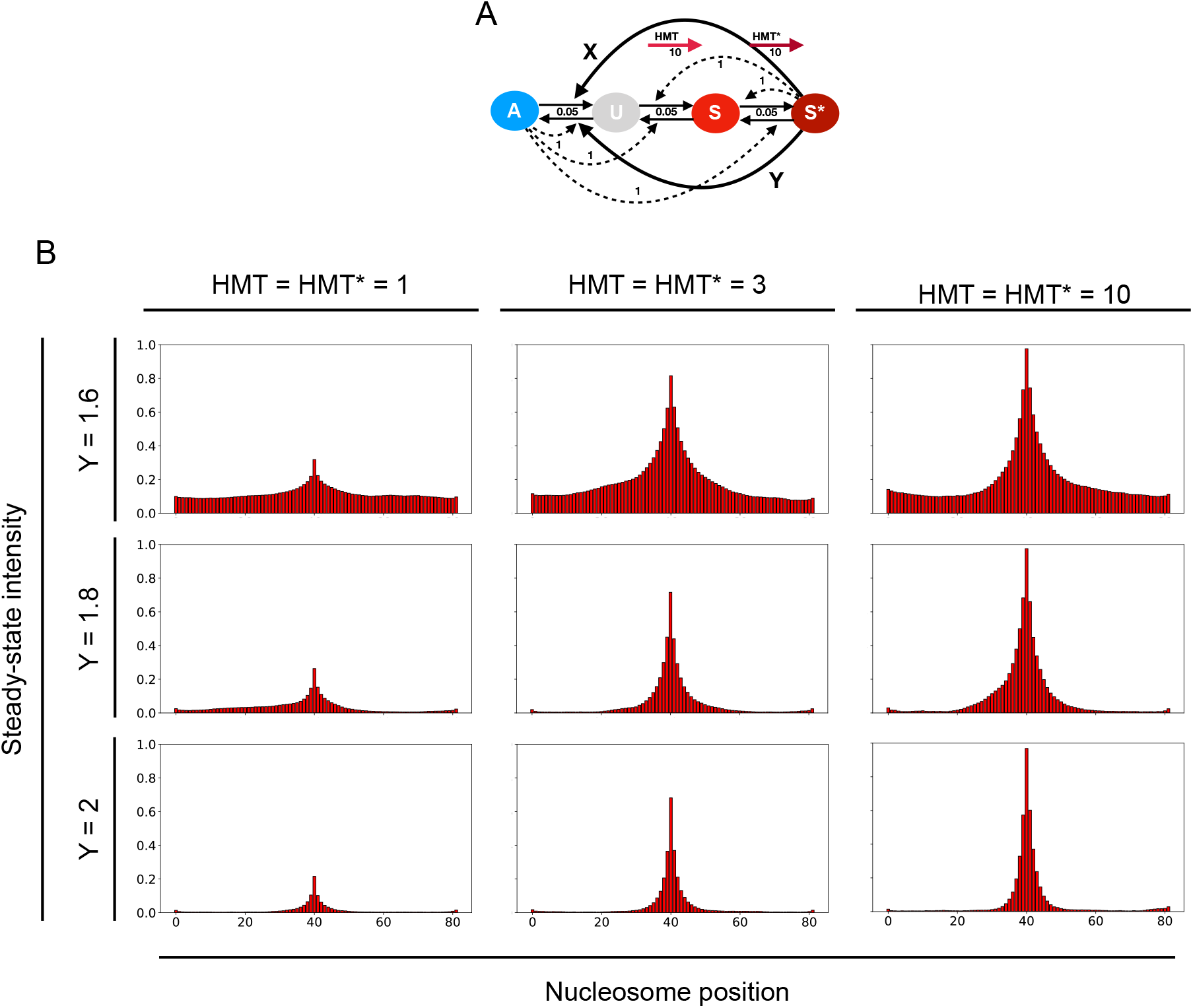
Localization of methylation around a silencer. A) Global feedback shown with solid lines depend on distance as 1/*d*. All local conversion rates are 1, *X* = 1.4, and HMT determines the strength of nucleation rates from central silencer. The direct conversion rates are all 0.05. B) Stregth of global negative feedback (Y) can localize the profile around a silencer. Nucleation strength (HMT and HMT*) mainly determines the height of the peak.

**Fig. S11.**
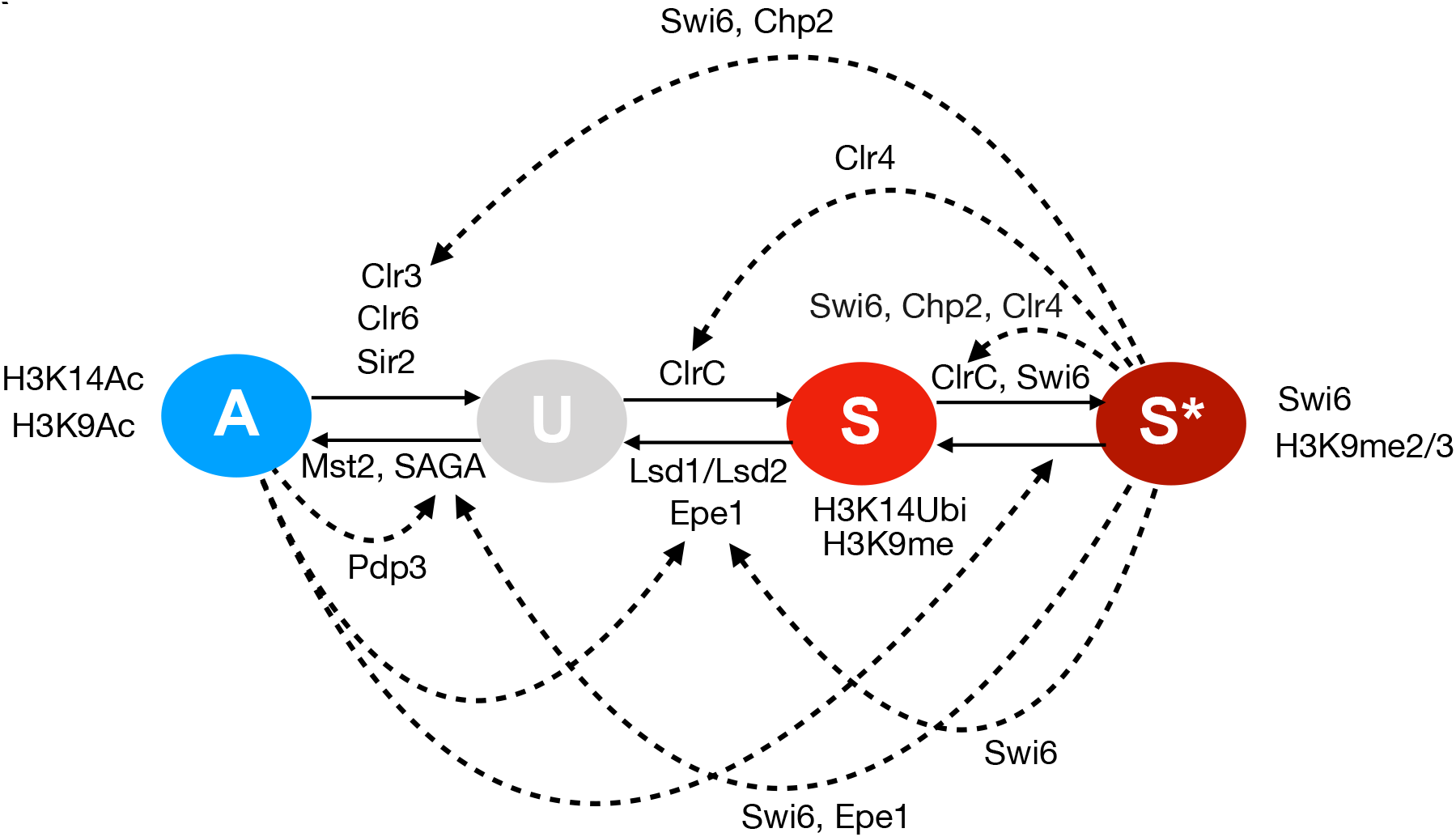
Schematics showing known interactions of read-write enzymes in *S Pombe* associated with the 4-state model. This schematic shows positive and negative feedback regarding the recruitment of read-write enzymes by specifically modified nucleosomes and their subsequent attempts to modify other nearby nucleosomes, e.g., through binding-dependent stimulation or disinhibition of their enzymatic activities. Swi6 and Chp2 are responsible for S*-mediated feedback th rough self-aggregation and binding to H3K9me nucleosomes (62) via its chromodomain, the recruitment of HDACs (26, 63, 64), including Clr3 (41, 65, 66) as well as a recruitment of the H3K9me transferase Clr4 (62). Another S*-mediated positive feedback occurs through Clr4, which recruits the Clrc complex responsible for H3K9me (20, 67, 68) via its Chromodomain (24, 69). Negative feedback potentially occurs through the S*mediated recruitment of Swi6, which associates with the putative H3K9me demethylase Epe1 (39) or with the SAGA complex (that contains an histone acetyl-transferase) via Epe1 1 (32). A-mediated positive feedback is less well characterized. Well established is that positive feedback A-mediated positive feedback occurs by recruiting the H3K14 acetyltransferase Mst2 via Pdp3 binding to A nucleosomes modified at H3K36me. Mst2 recruitment, in turn, stimulates H3K14 acetylation directly and indirectly through a positive feedback loop involving acetylation of the histone H2B ubiquitin ligase Brl1 by Mst2 (70). Positive feedback via A nucleosomes on S-to-U and S*-to-S conversions might occur through the Lsd1/Lsd2 demethylases (71). However, this remains to be investigated.

**Fig. S12.**
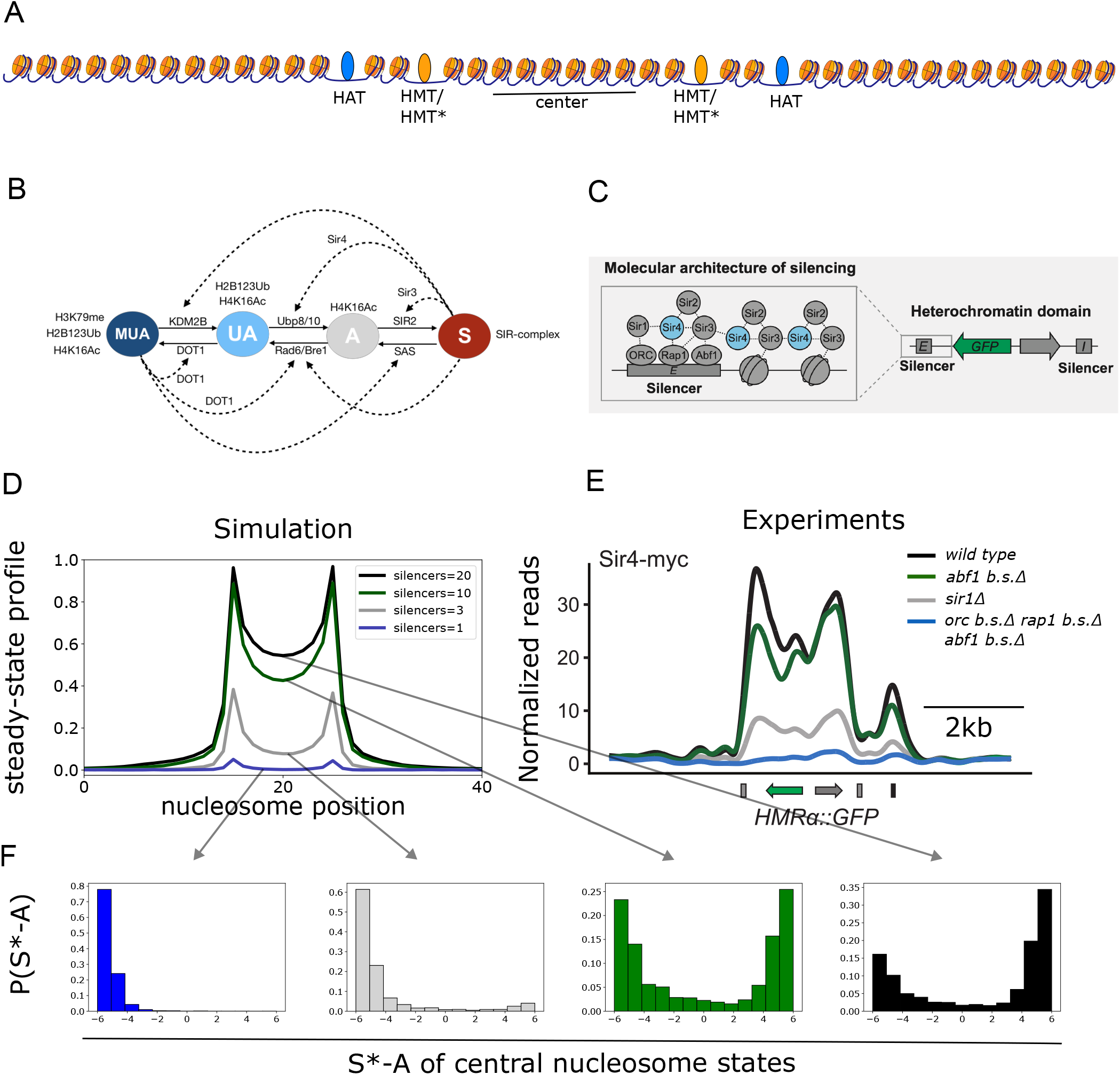
Nucleation strength controls the degree of silencing by modulating the switching rate between active and silent chromosomal states. A) Schematic of the region, showing only 40 nucleosomes, although simulations were made on 80 nucleosomes in total (20 additional nucleosomes on each side). Simulations were made with the same model and parameter values as in Fig. 6, except that an additional HAT activity (direct UtoA conversions) of 2 per update step has been performed on 2 nucleosomes at positions 33 and 47 (shown in A) and the nucleation strength has been adjusted. B) Overview of known interactions between nucleosomes and read-write enzymes. There is ample experimental evidence for extensive positive feedback towards the heterochromatic state in *budding yeast*. The SIR complex contains the H4K16 deacetylasease SIR2 (72) and SIR3 which binds to deacetylated nucleosomes more strongly than to acetylated ones and also reinforces further recruitment of SIR2/4 (72, 73). This interplay is thus believed to create a positive feedback/spreading of the SIR-complex from nucleation sites (74). Further, SIR4 interacts with the deubiquitinase Ubp10/Dot4p and might thus well be part of a positive feedback (75, 76). Moreover, there is indication that the just recently discovered H3K79me demethylase KDM2B in HEK294T cells functions in a SIRT1 dependent manner (77). However an H3K79me demethylase has not been found yet in budding yeast. Interestingly, positive feedback has been very recently suggested. E.g. (47) shows that H4K16Ac, H2B123Ub and H3K79me function together to stimulate the H3K79 methyltransferase activity of DOT1. Another positive feedback from the euchromatic site is suggested by (78) who show that DOT1 promotes ubiquitination by RAD6/Bre1. Evidence for a potential feedback on the acetyltransferase SAS and a potential negative feedback mediated by the SIR-complex remains to be shown. C Schematics adopted from (8) showing the different proteins involved in the recruitment of the SIR complex to the silencers and the spreading of the Sir complex to neighboring nucleosomes.. D) Steady-state profiles resulting from simulating the system with aforementioned parameter-values for 2 * 10^7^ steps (> 12.000 generations). E) normalized reads from ChIP-seq experiments of different S. *Cerevisiae* boundary mutants. F) Histograms showing the difference between the time-averaged number of S* and A state nucleosomes in the center corresponding to the differenr steady-state enrichments shown in D. Thus, the strength of the silencers determines the degree of silencing going from active to bistable to mostly silent without loosing confinement. This is in agreement with the recent findings in (8)

## Notes

### Competing Interest Statement

The authors have declared no competing interest.

### Summary of Updates

The revised paper changed Fig. S12 with correct references. Added reference to 2012 paper by Courtney Hodges and Gerald Crabtree

